# A regulator of amino acid catabolism controls *Acinetobacter baumannii* gut colonization

**DOI:** 10.1101/2025.09.11.674065

**Authors:** John H. Geary, Xiaomei Ren, Dziedzom A. Bansah, Ngoc T.T. Pham, Iván C. Acosta, Bryan M. Kigongo, Jonathan D. Winkelman, Francis Alonzo, Matthew T. Henke, Lauren D. Palmer

## Abstract

Asymptomatic gut colonization increases the risk of clinical infection and transmission by the multidrug-resistant pathogen *Acinetobacter baumannii*. Ornithine utilization was shown to be critical for *A. baumannii* competition with the resident microbiota to persist in gut colonization, but the regulatory mechanisms and cues are unknown. Here, we identify a transcriptional regulator, AstR, that specifically activates the expression of the *A. baumannii* ornithine utilization operon *astNOP.* Phylogenetic analysis suggests that AstR was co-opted from the *Acinetobacter* arginine utilization *ast(G)CADBE* locus and is specialized to regulate ornithine utilization in *A. baumannii*. Reporter assays showed that *astN* promoter expression was activated by ornithine but inhibited by glutamate and other preferred amino acids. *astN* promoter expression was similarly activated by incubation with fecal samples from conventional mice but not germ-free mice, suggesting AstR-dependent activation of the *astN* promoter responds to intermicrobial competition for amino acids. Finally, AstR was required for *A. baumannii* to colonize the gut in a mouse model. Together, these results suggest that pathogenic *Acinetobacter* species evolved AstR to regulate ornithine catabolism, which is required to compete with the microbiota during gut colonization.

## INTRODUCTION

*A. baumannii* is a common cause of hospital acquired infections and poses an urgent public health threat worldwide due to its widespread, extensive drug resistance^1,2^. *A. baumannii* can asymptomatically colonize any site in the human body and asymptomatic colonization is a direct risk factor for invasive infection^3,4^. While *A. baumannii* is not typically pathogenic in the gut, multiple reports highlight that gut colonization is an important reservoir for transmission of drug-resistant *A. baumannii*^5–13^. Gut colonization with *A. baumannii* can increase the risk of subsequent clinical infection by 2 to 15-fold^3,14–16^. While *A. baumannii* was rarely part of gut microbiome in healthy adults (<1% of participants)^17^, we recently reported that the prevalence in a cohort of healthy pre-weaning infants was 38 to 88%^18^. Furthermore, the prevalence in hospitalized individuals ranges from 15 to 41%^3,4,6,7,9,17,19,20^. Therefore, *A. baumannii* can colonize the human gut, which can result in transmission and increased risk of clinical infection in healthcare settings.

To colonize the gut, invading pathogens must respond to intermicrobial competition and niche exclusion by the resident microbiota. Competition for nutrients is considered a dominant mechanism of microbiota-mediated colonization resistance^21–24^. We recently showed that *A. baumannii* ornithine utilization is required to overcome microbiota competition and persistently colonize the gut in a mouse model^18^. However, it is not known how *A. baumannii* senses carbon sources and regulates ornithine catabolism to colonize the gut.

In this study, we investigated how *A. baumannii* regulates the ornithine catabolic operon. We previously demonstrated that the enzyme AstO is required for *A. baumannii* to utilize ornithine as a carbon and nitrogen source^18^. AstO is encoded in the *astNOP* operon, divergently transcribed from a gene encoding a predicted regulator that we named AstR^18^. AstR is annotated as an AsnC-family regulator, a subfamily of Feast-Famine Response Proteins (FFRP) exemplified by the leucine responsive protein Lrp^25,26^. In *E. coli,* the FFRP AsnC activates transcription of the divergently transcribed asparagine synthase gene (*asnA*): in low asparagine, the N-terminal DNA binding domain binds upstream of and activates *asnA*, upon binding asparagine with the C-terminal ligand binding domain activation ceases^27–29^. FFRP are encoded by nearly half of all sequenced bacteria and characterized examples detect small molecule metabolites and regulate the corresponding metabolic genes^25^.

Here, we show that the predicted AsnC-family regulator AstR is required for *A. baumannii* ornithine utilization, activation of the *astNOP* operon, and gut colonization in a mouse model. Together, these results suggest that AstR evolved to regulate *A. baumannii* ornithine catabolism, which is required for competing against the microbiota during gut colonization.

## RESULTS

### The predicted transcriptional regulator AstR is required for ornithine utilization in *A. baumannii* and arginine utilization in *A. baylyi*

The AST pathway catabolizes arginine and ornithine into glutamate in organisms such as *E. coli* and *P. aeruginosa*^30,31^ and we recently showed that AstO allows *A. baumannii* to utilize ornithine as the sole carbon and/or nitrogen source (Figure 1A)^18^. AstO is encoded in the *astNOP* operon, a partial duplication of the *ast* operon that is present in *A. baumannii* and other pathogenic *Acinetobacter* species (Figure 1B).^18^ The partially duplicated *ast* operon consists of *astN* (*astC* homolog), *astO* (*astA* homolog) and the putative amino acid transporter *astP*, divergently transcribed from a putative transcriptional regulator (Figure 1B). Δ*astO* cannot utilize ornithine as a sole carbon or nitrogen source but maintains the ability to utilize arginine^18^. However, it was not known how *A. baumannii* regulates the *astNOP* operon. A putative transcriptional regulator, which we named *astR,* is divergently transcribed from *astNOP* in the *Acinetobacter baumannii/calcoaceticus* complex containing all important pathogens in the genus (Figure 1B).

**Figure 1:**
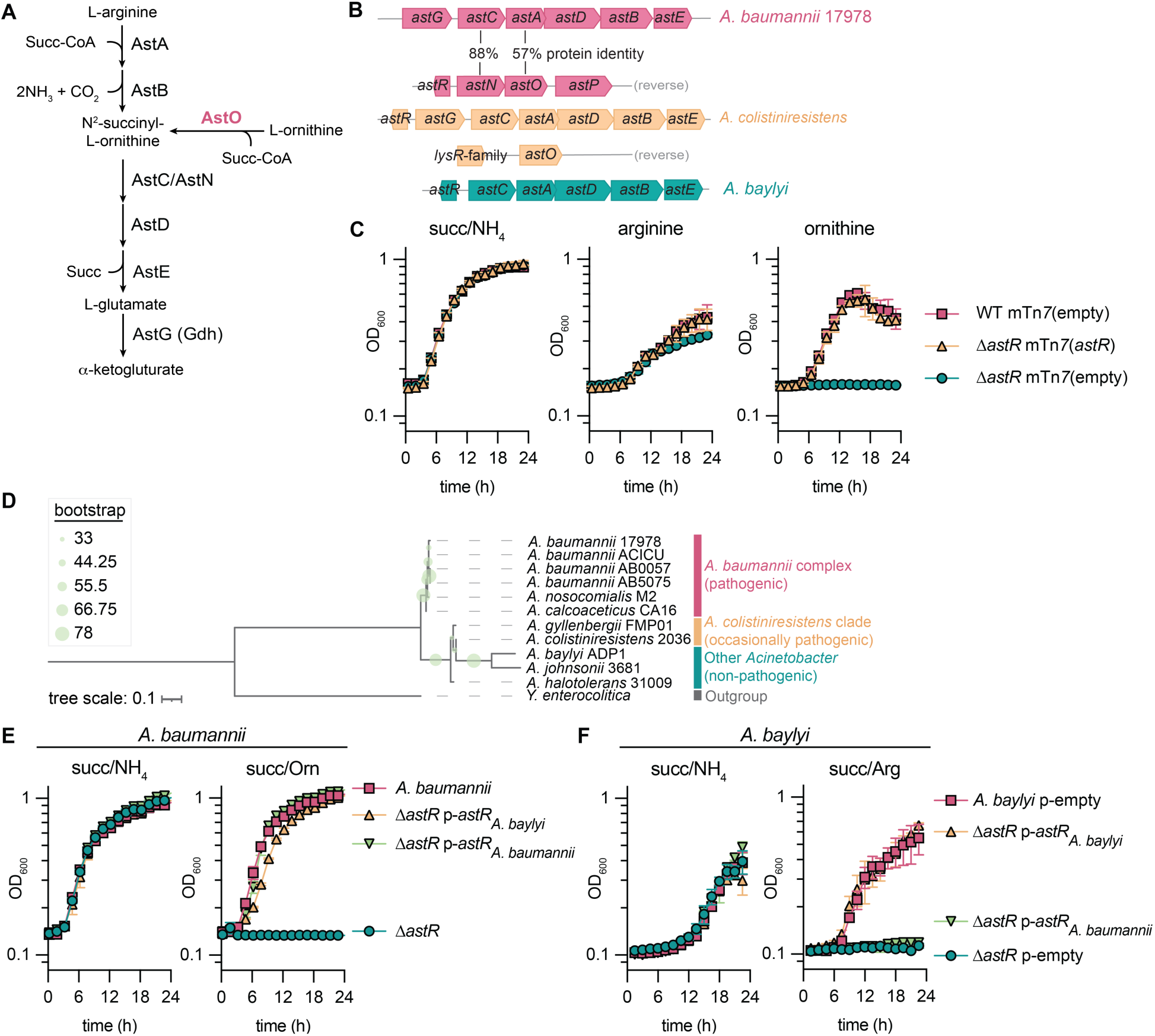
The predicted regulator AstR is required for ornithine and arginine catabolism in *Acinetobacter* species. **(A)** The arginine succinyltransferase (AST) arginine and ornithine utilization pathway in *A. baumannii*. **(B)** *ast* loci in *A. baumannii* ATCC 17978, *A. colistiniresistens*, and *A. baylyi*. **(C)** *A. baumannii* ATCC 17978VU wildtype (WT) and Δ*astR* strains with empty or *astR* complementation introduced by mini Tn*7* (mTn*7*) at the *att* site. Strains were grown in nitrogen-free M9 minimal media with succinate, arginine, or ornithine at 16.5 mM and NH_4_Cl at 18.6 mM, as indicated (n = 3; mean ± SD, experiments were performed at least twice with similar results). **(D)** Rooted phylogenetic tree of AstR protein sequences from representative *Acinetobacter* species with a *Yersinia enterocolitica* AsnC-family member as an outgroup. Tree scale in amino acid substitutions per site. **(E-F)** *A. baumannii* ATCC 17978VU or *A. baylyi* ATCC ADP1 wildtype and Δ*astR* strains with empty or *astR* complementation introduced in the pWH1266 plasmid. The pWH1266 plasmids used endogenous the *astR* promoter from the host strain. Strains were grown in nitrogen-free M9 minimal media with succinate at 16.5 mM and arginine, ornithine, or NH_4_Cl at 18.6 mM, as indicated (n = 3; mean ± SD, experiments were performed at least twice with similar results).

*astR* is only divergently encoded from the *astNOP* operon in the *Acinetobacter baumannii/calcoaceticus* complex (Figure 1B)^18^. In all other *Acinetobacter* spp., *astR* is divergently encoded from the full *ast* operon, typically *astGCADBE* (*astCADBE* in *Acinetobacter baylyi* ADP1). We hypothesized that *astR* encodes a transcriptional activator of the *astNOP* operon required for ornithine utilization in *A. baumannii*. An *A. baumannii* ATCC 17978 Δ*astR*::Kn (Δ*astR*) mutant grew similarly to wildtype (WT) with succinate as the sole carbon source and NH_4_ as the sole nitrogen source (Figure 1C). The Δ*astR* mutant also grew with arginine as the sole carbon and nitrogen source, but was unable to utilize ornithine (Figure 1C). There was a minor defect in stationary phase in the Δ*astR* mutant in arginine (Figure 1C), suggesting AstR may also sense arginine. These defects were complemented by reintroduction of the *astR* gene with its endogenous promoter at the *att* genomic locus by mini Tn*7* (mTn*7*) (Figure 1C). These data show AstR is required for ornithine utilization and suggest AstR may activate the *astNOP* operon in *A. baumannii*.

We next explored the phylogeny of AstR across *Acinetobacter* species. Using an AsnC-family member from *Y. enterocolitica* as an outgroup, an *Acinetobacter* AstR phylogenetic tree was constructed to explore the evolutionary relationships. AstR within the *Acinetobacter baumannii/calcoaceticus* complex clustered together while AstR outside of the *Acinetobacter baumannii/calcoaceticus* clade clustered together separately (Figure 1D). However, these proteins maintain high homology: AstR from *A. baumannii* ATCC 17978 and *A. baylyi* ATCC ADP1 have 69% identity and 88% similarity, with most predicted DNA binding and ligand binding residues conserved between the two (Figure S1A)^29^.

We next tested whether AstR from the nonpathogenic environmental species *A. baylyi* could complement *A. baumannii* Δ*astR*. A*. baylyi* does not encode AstO and cannot utilize ornithine but can utilize arginine as the sole nitrogen source.^18^ Thus, we hypothesized that *A. baylyi* AstR may be required for utilization of arginine as the sole nitrogen source, but could complement ornithine utilization in *A. baumannii* Δ*astR*. All *A. baumannii* strains grew with NH_4_ or arginine as the sole nitrogen source (Figure 1E, S1B). Similarly, all *A. baylyi* strains grew with NH_4_ and none grew with ornithine, as we previously reported (Figure 1F, S1C).^18^ *A. baumannii* Δ*astR* was unable to grow with ornithine as the sole nitrogen source, and this defect could be complemented by expression of *A. baumannii* or *A. baylyi astR* from a multi-copy plasmid (Figure 1E). For each complementation strain, the *astR* promoter (*astRp*) from the host species was used to drive *astR* expression. *A. baylyi* Δ*astR* was unable to grow with arginine as the sole nitrogen source as predicted (Figure 1F). For the *A. baylyi* Δ*astR* strain, complementation of *A. baylyi astR* restored growth with arginine, while complementation of *A. baumannii astR* did not (Figure 1F). This finding contrasted with results in *A. baumannii* Δ*astR* (Figure 1E) and suggests that *A. baumannii* AstR may be more specialized to regulate the *astNOP* operon. Together, these results show that AstR is conserved throughout *Acinetobacter* and required for arginine utilization in *A. baylyi* and ornithine utilization in *A. baumannii*, suggesting it may be important for activation of ornithine utilization for *A. baumannii* to colonize the host.

### AstR is required for activation of *astNOP* operon expression

To test whether *A. baumannii* AstR may activate expression of *astNOP* operon, a luciferase reporter plasmid was employed (p-*astNp*-*lux*). First, there was little expression from the *astNp* promoter in minimal media with succinate or ornithine as the sole carbon source and there was no growth (n.g.) for Δ*astR* in the ornithine/NH_4_ medium (Figure 2A, S2A). Since the Δ*astR* mutant cannot utilize ornithine, strains were grown in succinate/NH_4_ with ornithine (16.5 mM) to test the effect of ornithine on the *astNp*-lux reporter in the Δ*astR* mutant. The expression from the *astNp* promoter was dramatically increased when WT was grown with succinate/ornithine/NH_4_ (Figure 2A, S2A). However, the Δ*astR* strain was unable to activate *astNp* in ornithine/succinate/NH_4_ medium and luminescence was significantly lower (Figure 2A, S2A). Arginine also promoted AstR-dependent activation of expression from the *astNp* operon (Figure 2A, S2A), suggesting that *A. baumannii* AstR maintains the ability to sense arginine. The *astR* promoter (*astRp*) was similarly expressed in the conditions tested in the WT or Δ*astR* strain background (Figure S2B), suggesting *astR* is not autogenously regulated, in contrast to *E. coli* AsnC^27^.

**Figure 2:**
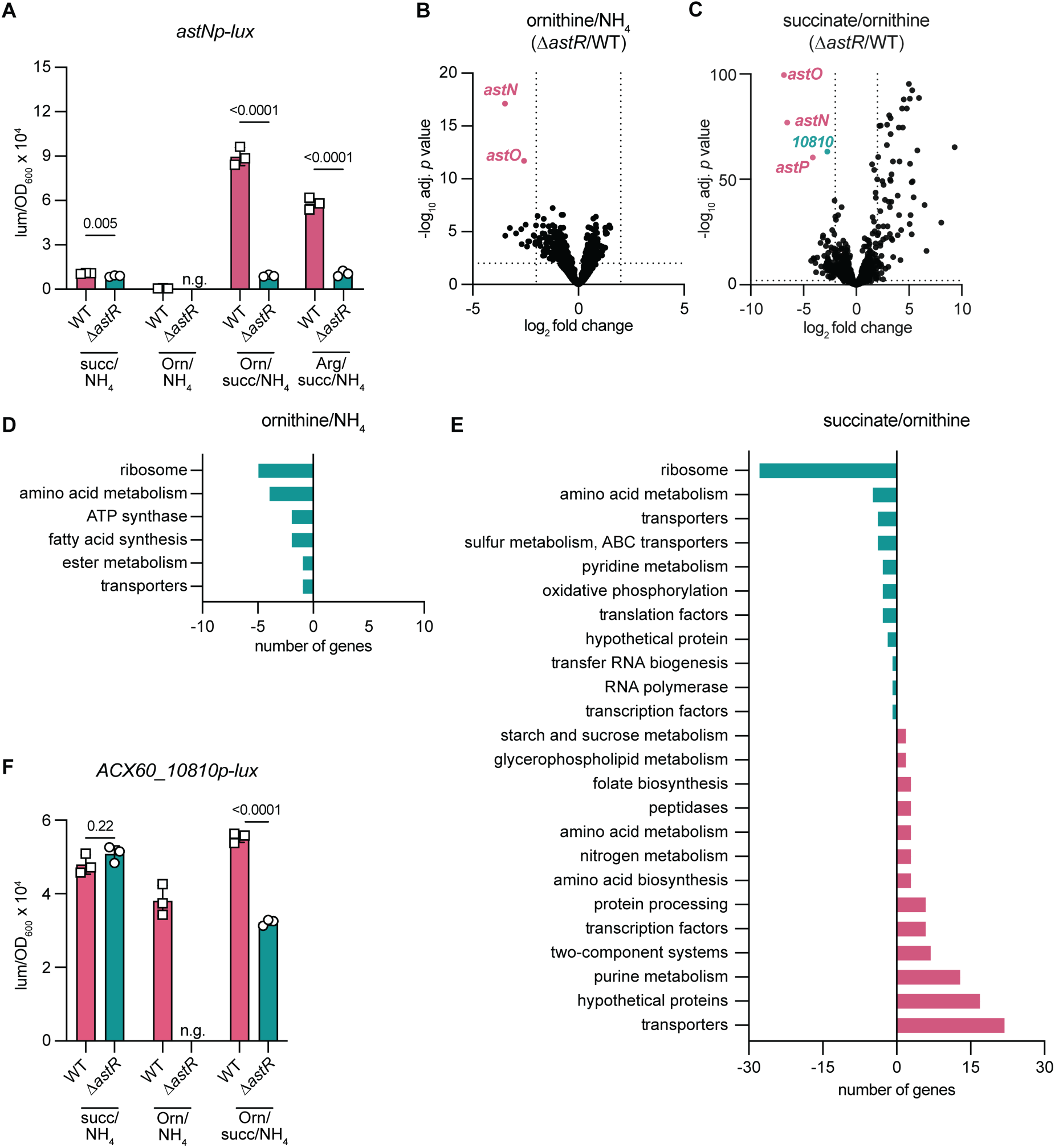
*A. baumannii* AstR is required for activation of the *astNOP* ornithine catabolic operon. **(A)** *A. baumannii* wildtype (WT) and Δ*astR* strains containing an *astN* promoter (*astNp*) driving luciferase expression from the *lux* operon were grown in nitrogen-free M9 minimal media minimal media with succinate (succ), ornithine (Orn), and arginine at 16.5 mM and NH_4_Cl at 18.5 mM, as indicated. Luminescence was normalized to optical density at 600 nm (OD_600_) at 0.4. N.g. indicates no growth in that condition (n = 3; mean ± SD; *p* by multiple unpaired *t*-tests; experiments were performed at least twice with similar results). **(B-C)** Volcano plot of Δ*astR*/WT after 1 h exposure to nitrogen-free M9 minimal medium with ornithine at 16.5 mM and NH_4_Cl at 18.6 mM (B) or succinate at 16.5 mM and ornithine at 18.6 mM (C). Each dot is one gene (n = 3; adj. *p* value and fold-change were determined by DESeq2; dotted lines shown at |log2 fold change|=2 and adj. *p* value=0.05). **(D-E)** KEGG pathway analysis of the number of genes with significantly changed expression in ornithine/NH_4_ M9 minimal media shown in panel B or succinate/ornithine M9 minimal media shown in panel C. Genes with decreased expression shown on the negative x-axis and with increased expression on the positive x-axis. Significance was |log_2_fold-change| ≥ 2, adjusted *p* value <0.05. **(F)** *A. baumannii* WT and Δ*astR* strains containing an *ACX60_10810* promoter (*ACX60_10810p*) driving luciferase expression from the *lux* operon were grown in nitrogen-free M9 minimal media minimal media with succinate and ornithine at 16.5 mM and NH_4_Cl at 18.5 mM, as indicated. Luminescence was normalized to OD_600_ at 0.4. n.g. indicates no growth in that condition (n = 3; mean ± SD, *p* by multiple unpaired *t*-tests; experiments were performed at least twice with similar results). *astNp, astNOP* operon promoter; lum, luminescence; OD_600_, optical density at 600 nm; WT, wildtype; succ, succinate; Orn, ornithine, n.g., no growth; adj. *p* value, adjusted *p* value; *10810*, *ACX60_10810*; *ACX60_10810p*, *ACX60_10810* promoter.

To characterize the AstR regulon, RNA sequencing was performed on *A. baumannii* ATCC 17978 WT and Δ*astR* when incubated in minimal media with ornithine as the sole carbon or sole nitrogen source for 1 h to capture initial transcriptional responses. With incubation in ornithine/NH_4_ where ornithine is the sole carbon source, *astN* and *astO* were significantly downregulated in Δ*astR* compared to WT (Figure 2B). When incubated with ornithine as the sole nitrogen source (succinate/ornithine), all genes in the *astNOP* operon were significantly downregulated in Δ*astR* compared to WT (Figure 2C). The fact that *astP* expression was only changed in the succinate/ornithine condition suggests that there may be additional regulation in the 399 bp intergenic region between *astO* and *astP*. Published RNA-seq data suggest that *astP* is co-transcribed with *astNO*^32^; additionally, an *astP* promoter reporter with the 399 bp *astO-astP* intergenic region had no activity in any medium tested including succinate/ornithine/NH_4_ (Figure S2C). Thus, additional regulation of *astP* may be post-transcriptional.

Next, expression of the *ast* operons was compared. The fold change for *astN* and *astO* in the WT RNA-seq data was almost 16-times higher in ornithine as sole nitrogen source (succinate/Orn) than ornithine as sole carbon source (Orn/NH_4_; Figure 2B, S2D). This fold change is consistent with the difference in the *astNp-*driven luminescence with ornithine and with or without the addition of succinate (Figure 2A). Additionally, the *astGCADBE* operon was upregulated in WT in succinate/Orn compared to Orn/NH_4_ (Figure S2D), consistent with the conserved role of the AST pathway in utilization of arginine and ornithine as sole nitrogen sources throughout the *Acinetobacter* genus. Comparison of WT to Δ*astR* confirmed that in *A. baumannii* ATCC 17978, AstR does not appear to regulate the *astGCADBE* operon (Figure 2B-C).

Pathway analysis was performed on genes that were significantly downregulated or upregulated in Δ*astR* compared to WT. In ornithine/NH_4_, downregulated genes in Δ*astR* were predominantly in the ribosome or amino acid metabolism pathways, and there were no significantly upregulated genes in *ΔastR* (Figure 2D). In succinate/ornithine, downregulated genes in Δ*astR* were also predominantly in ribosome or amino acid metabolism pathways, and upregulated genes were in purine metabolism, hypothetical proteins, and transporters (Figure 2E). Because Δ*astR* cannot utilize ornithine as the sole carbon or nitrogen source, some gene expression changes may be due to inability to grow in these conditions.

The third most downregulated pathway in Δ*astR* in succinate/ornithine was transporters, including the gene *ACX60_10810*, which is annotated as encoding a putative dicarboxylate symporter. *ACX60_10810* was downregulated in succinate/ornithine in a similar trend to *astNOP* genes in *ΔastR* (Figure 2C). Using a luciferase reporter, expression from the *ACX60_10810* promoter (*10810p*) was high in all conditions where strains grew and was significantly higher in WT than Δ*astR* when grown in ornithine/succinate/NH_4_ (Figure 2F, S2F). This suggests that *ACX60_10810* has high baseline expression in these conditions and AstR may further activate *10810p* in the presence of ornithine. However, this effect could be indirect due to other metabolic differences from the inability of Δ*astR* to catabolize ornithine. Together, these results suggest that AstR activates the *astNOP* operon and *ACX60_10810*.

### AstR binds the *astNOP* promoter

AsnC-family transcriptional regulators typically function by directly binding to a DNA promoter region to alter expression^25,26^. Electrophoretic mobility shift assays (EMSA) were used to assess interaction of purified AstR-His_6_ incubated with Cy5.5-labeled *astNp* (135 bp region, Figure 3A). First, His_6_-AstR was confirmed to function by complementation of the *ΔastR* strain in minimal ornithine medium (Figure S3A). By EMSA, AstR bound the *astNp* in a concentration-dependent manner (Figure 3B). The gel shift was confirmed to be due to His_6_-AstR binding by supershifting the bound DNA with an anti-His antibody (Figure S3B). Next, *astNp* truncations were assessed to determine the region of AstR binding. A region >100 bp upstream of the *astN* translation start site was required for AstR binding by EMSA (Figure 3C). Similarly, this region 100-135 bp upstream of the *astNp* translation start site was required for activation in succinate/ornithine/NH_4_ (Figure 3D, S3C). These data suggest the region 100-135 bp upstream of the *astNp* translation start site contains a binding site for AstR-dependent activation.

**Figure 3:**
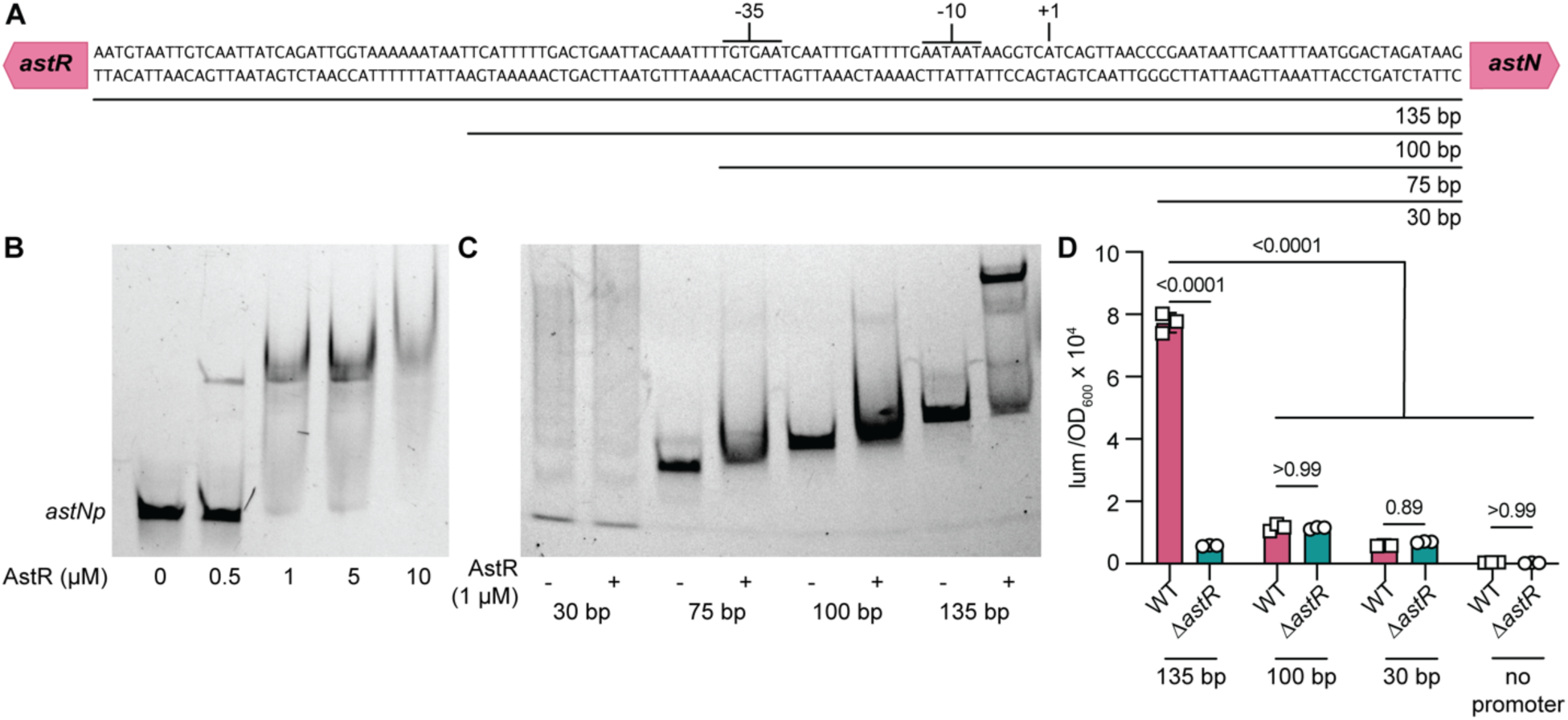
AstR directly binds *astNOP* promoter DNA. **(A)** The intergenic region between *astR* and *astN* is depicted including the predicted *astR* -35 and -10 RNA polymerase binding sites and +1 transcription start site. **(B)** The 135 bp intergenic region between *astR* and *astN* was amplified with a Cy5.5-labeled oligonucleotide. Electrophoretic mobility shift assays (EMSA) were performed with purified His_6_-AstR at the indicated concentrations. Results are representative of at least 2 independent experiments. **(C)** The 30 bp, 75 bp, 100 bp, or 135 bp intergenic regions upstream of *astN* as shown in panel A were amplified with a Cy5.5-labeled oligonucleotide and incubated with purified His6-AstR for EMSA. **(D)** The 135 bp, 100 bp, and 30 bp intergenic regions upstream of *astN* as shown in panel A were cloned upstream of a *lux* luciferase reporter or with a no promoter control. WT and Δ*astR* strains with reporters were incubated in nitrogen-free M9 minimal medium with succinate and ornithine at 16.5 mM and NH_4_Cl at 18.6 mM. Luminescence was normalized to optical density at 600 nm (OD_600_) at 0.4 (n=3; *p* by two-way ANOVA with Sidak’s multiple comparisons). *astNp, astNOP* operon promoter; lum, luminescence; OD_600_, optical density at 600 nm.

AsnC-family regulators often oligomerize and regulate gene expression in response to ligand binding^25,26^. However, addition of ligands such as arginine, ornithine, or arginine and succinate together did not dramatically alter His_6_-AstR binding to *astNp* DNA by EMSA (Figure S3D-F). We next investigated the oligomeric and ligand-binding states of the purified protein. The predicted His_6_-AstR monomer molecular weight is 18.1 kDa; size exclusion chromatography with standard curves estimated that the His_6_-AstR was 31.8 kDa (Figure S3G-I), suggesting it is a dimer. Native and denatured mass spectrometry showed that His_6_-AstR purified with modifications typical of expression in *E. coli* but with no ligand bound (Figure S4). Thus, these data show that dimeric apo-AstR directly binds the *astNOP* promoter *in vitro* and that binding and AstR-dependent activation requires a region 100-135 bp upstream of the *astNOP* promoter.

### Preferred amino acid carbon sources inhibit *astNOP* operon expression

Our previous study showed that ornithine is not a preferred carbon source for *A. baumannii*. Rather, glutamate/glutamine/asparagine/histidine are preferred over succinate, which is preferred over arginine/ornithine.^18^ We hypothesized that the addition of preferred carbon sources may inhibit expression from the *astN* promoter. In succinate/ornithine/NH_4_ media that supports high expression from *astNp*, addition of preferred carbon sources glutamate, glutamine, asparagine, aspartate, and histidine led to significantly reduced *astNp* expression, while arginine did not affect expression (Figure 4A, S5A). Using glutamate, the inhibition of expression from the *astNp* operon was concentration dependent (Figure 4B, S5B). However, addition of glutamate or α-ketoglutarate, the product of glutamate degradation from the AST pathway, did not alter *astNp* DNA binding by purified His_6_-AstR, suggesting these effects could be indirect (Figure S5C). These data show expression from the *astNp* promoter is inhibited by addition of preferred amino acid carbon sources in the growth media.

**Figure 4:**
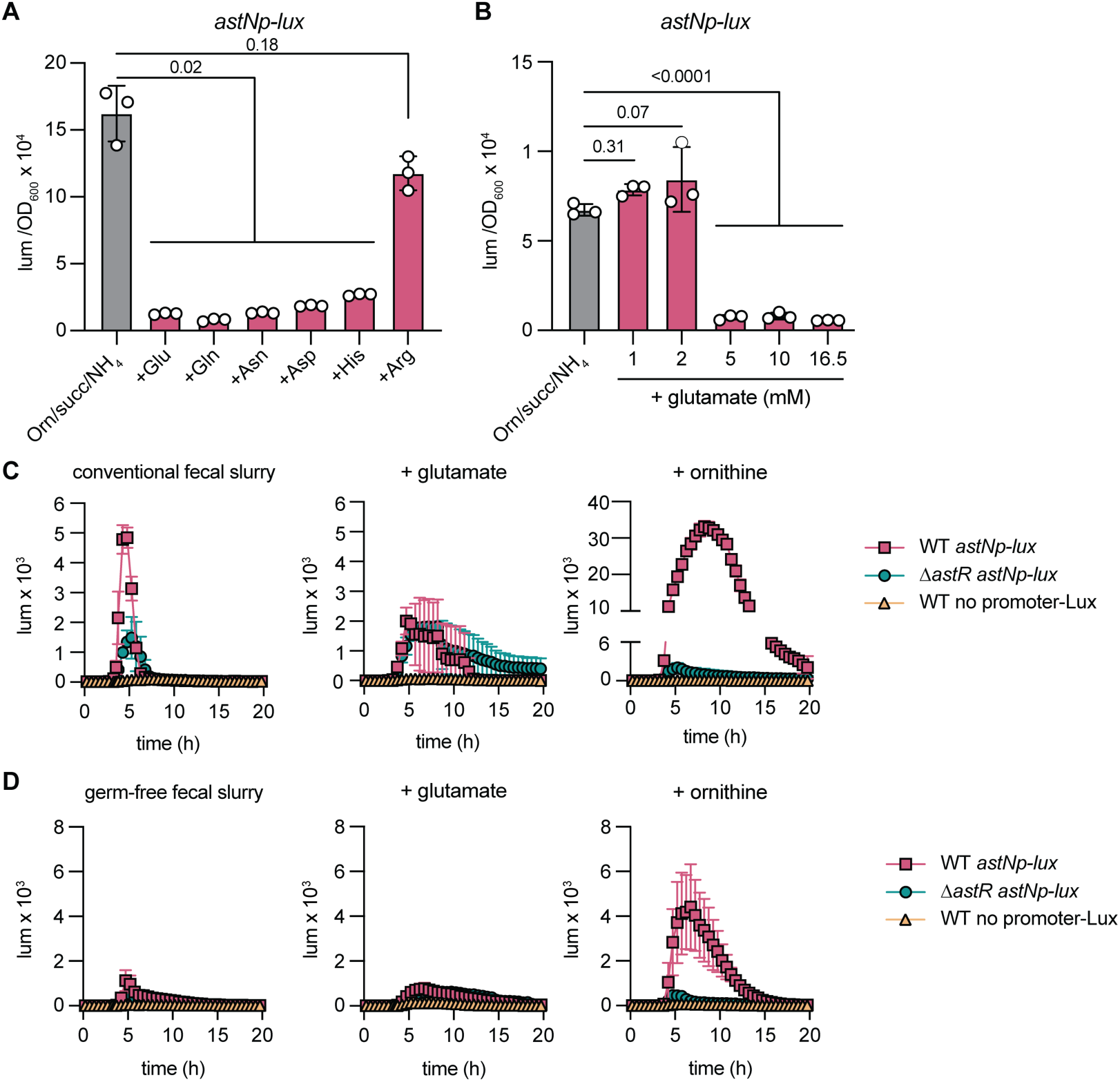
Preferred amino acid carbon sources inhibit AstR-dependent activation of the *astNOP* ornithine catabolic operon. **(A-B)** *A. baumannii* wildtype containing an *astN* promoter (*astNp*) driving luciferase expression from the *lux* operon was grown in nitrogen-free M9 minimal media minimal media with succinate (succ) and ornithine (Orn) at 16.5 mM and NH_4_Cl at 18.5 mM. In (A), 16.5 mM additional amino acids were added as indicated. In (B), glutamate was added at the indicated concentrations. Luminescence was normalized to optical density at 600 nm (OD_600_) at 0.4 (n=3; mean ± SD; *p* by one-way ANOVA with Dunnett’s multiple comparisons to no additional amino acids; experiments were performed at least twice with similar results). **(C-D)** Luminescence reporter curves for the indicated strains in fecal samples from conventional mice or germ-free mice homogenized in phosphate buffered saline (PBS). Additional glutamate or ornithine were added at 16.5 mM as indicated (n = 3; mean ± SD). *astNp, astNOP* operon promoter; lum, luminescence; Orn, ornithine; WT, wildtype; OD_600_, optical density at 600 nm.

We previously showed that *A. baumannii* requires ornithine catabolism to persistently colonize the gut of conventional mice, but that addition of preferred amino acid carbon sources such as glutamate rescue the Δ*astO* mutant^18^. This suggested that glutamate may inhibit expression from *astNp* in the gut microbiota environment. To test whether the *astNOP* operon is expressed in the gut, we examined the *astNp-lux* expression in fecal sample slurries from untreated conventional mice, a physiologically relevant *ex vivo* environment^33,34^. In fecal slurries from conventional mice, AstR-dependent expression from the *astN* promoter was activated (Figure 4C). Addition of glutamate inhibited this activation, while ornithine promoted AstR-dependent expression from *astNp* (Figure 4C). These data suggest that in the gut environment with an intact microbiota present, AstR activates expression from the *astN* promoter.

However, in germ-free mice that lack a microbiota, we previously reported that the Δ*astO* mutant had no defect, likely due to lack of competition for preferred amino acid carbon sources^18^. Thus, we predicted that expression from the *astNp* operon would not be activated in fecal slurries from germ-free mice. Consistent with this model, there was little luminescence from the *astNp*-*lux* reporter in fecal slurries from germ-free mice without or with additional glutamate (Figure 4D). Additional ornithine promoted AstR-dependent activation of the *astNp*-lux reporter to a level similar to in fecal slurries from conventional mice, suggesting additional ornithine could overcome inhibition by available preferred amino acid carbon sources (Figure 4D). Together, these data show that AstR is required for activation of the *astN* operon in an *ex vivo* model of competition for preferred amino acids with the gut microbiota.

### AstR promotes gut colonization in mice

Because ornithine catabolism is required for gut colonization and the data thus far show AstR is required for ornithine catabolism, we predicted that AstR would be required for gut colonization in conventional mice. Using a post-antibiotics model, female Swiss-Webster mice were orogastrically inoculated with a 1:1 mixture of *A. baumannii* ATCC 17978 WT:Δ*astR* (Figure 5A). In this model, significantly more *A. baumannii* WT than Δ*astR* were shed in feces at 9 days post inoculation (Figure 5B). At 10 days, in the small intestine, cecum, and colon, *A. baumannii* WT were detectable and had significantly higher burdens than Δ*astR*, which was below the limit of detection (LOD) (Figure 5C). Competition in a fecal slurry model showed Δ*astO* and Δ*astR* had similar defects compared to WT *A. baumannii* in fecal samples from male and female mice (Figure S6). Together, these data show that AstR is necessary for persistent gut colonization in conventional mice.

**Figure 5:**
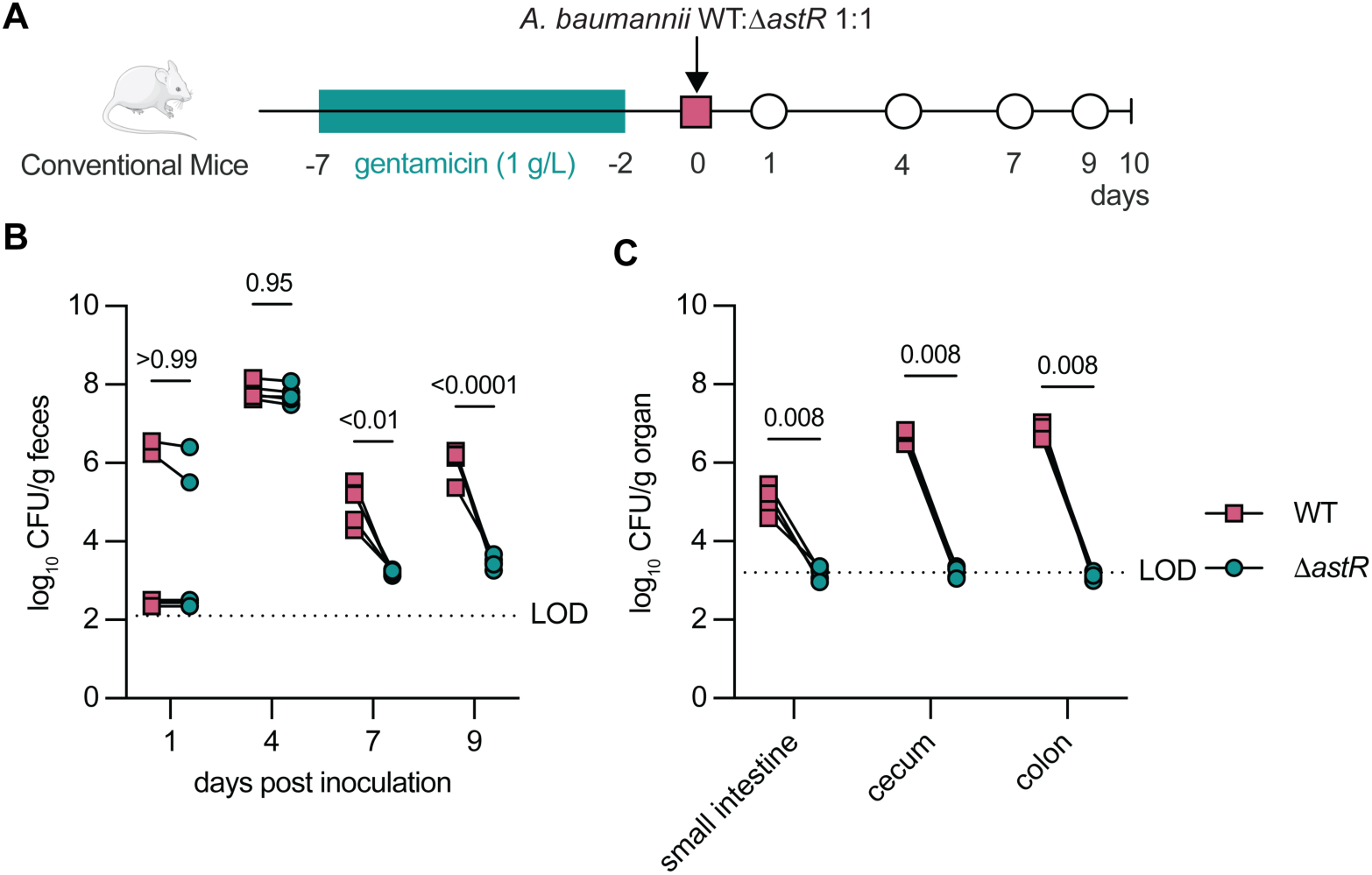
AstR is required for persistent gut colonization. **(A)** Experimental design of post-gentamicin *A. baumannii* gut colonization model. **(B)** *A. baumannii* wildtype (WT) and g*astR* fecal shedding over time (n=5; *p* by 2-way ANOVA with Sidak’s multiple comparisons). **(C)** *A. baumannii* WT and Δ*astR* enumerated from the indicated tissues collected at 10 days post inoculation (n=5; *p* by Mann-Whitney test). Lines connect colony-forming units (CFU) enumerated from the same mouse. WT, wildtype; CFU, colony-forming units; LOD, approximate limit of detection.

## DISCUSSION

Little is known about the molecular mechanisms *A. baumannii* uses to colonize and persist in the gut, a critical reservoir in healthcare settings. We previously reported that *A. baumannii* ornithine catabolism is required for persistent gut colonization in a mouse model^18^. However, how *A. baumannii* regulates ornithine catabolism remained unclear. Here, we characterize the *Acinetobacter* transcriptional regulator AstR that is required to activate the ornithine catabolic operon in *A. baumannii* and utilize ornithine as the sole carbon or nitrogen source.

AstR is an AsnC-family regulator that is specific to the *Acinetobacter* genus. Outside of the *A. baumannii/calcoaceticus* complex that includes all major pathogens, most *Acinetobacter* species use arginine and/or ornithine as nitrogen sources but not carbon sources^35,36,18^. Data included here show enhanced AstR-dependent activation of the *astNOP* operon in the presence of the organic acid succinate. Together, these findings suggests that AstR may have evolved in the *Acinetobacter* genus to activate the AST pathway to utilize arginine as a nitrogen source. The genomic organization, protein similarity, response to arginine, and potential for *A. baylyi astR* to cross-complement *A. baumannii* Δ*astR* suggest that *A. baumannii* AstR was co-opted from the *ast(G)CADBE* operon to regulate ornithine utilization as a carbon and nitrogen source. Results in this study show that *A. baumannii* AstR is required to activate expression from the *astN* promoter in the gut metabolite environment from conventional but not germ-free mice in *ex vivo* samples. Furthermore, AstR is required for persistent gut colonization in mice. Together, these data suggest that *A. baumannii* AstR activates ornithine catabolism to compete with the microbiota and persist in the gut.

Data included here demonstrate that His_6_-AstR can complement Δ*astR* and that it purifies as a dimer in apo-form that can bind the *astNOP* promoter. Additionally, glutamate and other preferred amino acid carbon sources inhibited expression from the *astNOP* promoter. However, addition of arginine, ornithine, succinate, or glutamate did not dramatically change *astNp* DNA binding by AstR by EMSA. Thus, how AstR integrates these nutrient signals to regulate the *astNOP* operon remains unclear and may involve additional transcriptional and/or post-transcriptional regulators. For example, a recent *A. baumannii* study showed that disruption of the two-component system response regulator gene *gacA* prevented activation of the *astGCADBE* operon and growth on arginine or ornithine^37^. It is not known whether GacA also regulates the *astNOP* operon and if its regulation of the *astGCADBE* operon is direct. Future studies could also investigate the potential for ligand binding to affect higher order oligomerization of AstR dimers and the effect on transcriptional regulation, which was recently reported for the FFRP Lrp in *E. coli*^38^. Nevertheless, the data here collectively suggest that the *astNOP* operon is maximally activated in an AstR-dependent mechanism to utilize arginine or ornithine as nitrogen sources.

Arginine and ornithine are emerging as key signals for bacterial interactions in the host^39^. For example, arginine sensing by ArgR modulates the virulence of enterohemorrhagic *Escherichia coli*, *Citrobacter rodentium*, *Salmonella enterica*, and hypervirulent *Klebsiella pneumoniae*^40–42^. Conversely, *Clostridioides difficile* and *Clostridium botulinum* increase toxin production in response to arginine depletion^43,44^. Arginine can also be degraded to ornithine by the host and microbiota^43,45–47^. Ornithine plays an important role in microbiota interactions in the gut, for example ornithine produced by *Lactobacilli* maintains a healthy gut mucosa^45^. For *C. difficile*, ornithine promotes asymptomatic colonization, toxin production, and spore formation^48–50^. Thus, arginine and ornithine metabolism and signaling likely play important roles for persistent gut carriage by invading pathogens.

Collectively, this study shows that the *A. baumannii* transcriptional regulator AstR is required for ornithine catabolism, activation of the *astNOP* ornithine catabolism operon, and gut colonization. The data presented here suggest that *A. baumannii* AstR was co-opted from the *Acinetobacter ast(G)CADBE* arginine catabolic locus and evolved to regulate ornithine catabolism in *A. baumannii* and other members of the *Acinetobacter baumannii/calcoaceticus* complex. This study adds to a growing body of evidence that arginine and ornithine are important signals and nutrient sources for pathogens in the gut. Better understanding how antimicrobial resistant pathogens such as *A. baumannii* sense and regulate nutrient metabolism during gut colonization will lay the foundation for developing treatment strategies to decolonize high-risk patients and prevent infections.

## MATERIALS AND METHODS

### Bacterial strains and growth

All strains generated for this manuscript are derived from *A. baumannii* ATCC 17978VU,^51^ unless otherwise noted (Table 1). Cloning was performed in *E. coli* DH5α λ*pir116*. Overnight cultures were grown starting from a single colony in Miller lysogeny broth (LB) at 37°C with shaking for 8-16 h. Solid media contained 1.5% w/v agar. Antibiotics were used in the following concentrations: carbenicillin 75 mg/L; kanamycin, 40 mg/L; chloramphenicol, 15 mg/L; sulfamethoxazole, 100 mg/L.

**Table 1:** Strains.

| Strain | Source | Identifier |
| --- | --- | --- |
| *Acinetobacter baumannii*<br>ATCC 17978VU (wildtype;<br>WT) | (Wijers *et al.*, 2021)<sup>51</sup> | LP486 |
| *Acinetobacter baumannii*<br>ATCC 17978VU $\Delta astO::Kn$ | (Ren and Clark, *et al.*, 2025)<sup>18</sup> | LP814 |
| *Acinetobacter baumannii*<br>ATCC 17978VU $\Delta astR::Kn$ | This manuscript | LP844 |
| *Escherichia coli*Bl21(DE3)<br>RIL | Agilent (Santa Clara,<br>California) | ArcticExpress (DE3) RIL<br>Competent Cells |
| *Escherichia coli*Bl21(DE3)<br>RIL pET15b-*his6-astR* | This manuscript | LP819 |
| *Acinetobacter baumannii*<br>ATCC 17978VU (wildtype;<br>WT) mTn7(empty) | (Ren and Clark, *et al.*, 2025)<sup>18</sup> | LP526 |
| *Acinetobacter baumannii*<br>ATCC 17978VU $\Delta astR::Kn$<br>mTn7(empty) | This manuscript | LP989 |
| *Acinetobacter baumannii*<br>ATCC 17978VU $\Delta astR::Kn$<br>mTn7(*astR*) | This manuscript | LP1165 |
| *Acinetobacter baumannii*<br>ATCC 17978VU pWH1266 | This manuscript | LP1178 |
| *Acinetobacter baumannii*<br>ATCC 17978VU $\Delta astR::Kn$<br>pWH1266 | This manuscript | LP907 |
| *Acinetobacter baumannii*<br>ATCC 17978VU $\Delta astR::Kn$<br>pLDP337(pWH1266-*astRpA*.<br>*baumannii-astRA. baumannii*) | This manuscript | LP1173 |
| *Acinetobacter baumannii* ATCC 17978VU $\Delta astR::Kn$ pLDP345(pWH1266-*astRpA*.<br>*baumannii-astR*<sub>A. baylyi</sub>) | This manuscript | LP1219 |
| *Acinetobacter baylyi* ATCC ADP1 | ATCC | LP460 |
| *Acinetobacter baylyi* ATCC ADP1 $\Delta astR::Kn$ | This manuscript | LP1084 |
| *Acinetobacter baylyi* ATCC ADP1 pWH1266 | This manuscript | LP1174 |
| *Acinetobacter baylyi* ATCC ADP1 $\Delta astR::Kn$ pWH1266 | This manuscript | LP1175 |
| *Acinetobacter baylyi* ATCC ADP1 $\Delta astR::Kn$ pLDP336(pWH1266-*astRpA*.<br>*baylyi-astR*<sub>A. baylyi</sub>) | This manuscript | LP1176 |
| *Acinetobacter baylyi* ATCC ADP1 $\Delta astR::Kn$ pLDP344(pWH1266-*astRpA*.<br>*baylyi-astR*<sub>A. baumannii</sub>) | This manuscript | LP1220 |
| *Acinetobacter baumannii* ATCC 17978VU (wildtype; WT) mTn7(empty) pLDP92(pLDP87-*astNp-lux*) | This manuscript | LP542 |
| *Acinetobacter baumannii* ATCC 17978VU $\Delta astR::Kn$ mTn7 (empty) pLDP92(pLDP87-*astNp-lux*) | This manuscript | LP998 |
| *Acinetobacter baumannii* ATCC 17978VU $\Delta astR::Kn$ mTn7 (*astR*) pLDP92(pLDP87-*astNp-lux*) | This manuscript | LP543 |
| *Acinetobacter baumannii* ATCC 17978VU mTn7(empty) pLDP108(*astRp-lux*) | This manuscript | LP605 |
| *Acinetobacter baumannii*<br>ATCC 17978VU $\Delta astR::Kn$<br>mTn7(empty)<br>pLDP108(*astRp-lux*) | This manuscript | LP606 |
| *Acinetobacter baumannii*<br>ATCC 17978VU $\Delta astR::Kn$<br>pLDP289(*ACX60_10810p-lux*) | This manuscript | LP1022 |
| *Acinetobacter baumannii*<br>ATCC 17978VU<br>pLDP289(*ACX60_10810p-lux*) | This manuscript | LP1023 |
| *Acinetobacter baumannii*<br>ATCC 17978VU<br>pLDP146(*astPp-lux*) | This manuscript | LP718 |
| *Acinetobacter baumannii*<br>ATCC 17978VU $\Delta astR::Kn$<br>pLDP146(*astPp-lux*) | This manuscript | LP719 |
| *Acinetobacter baumannii*<br>17978VU no promoter-*lux* | This manuscript | LP465 |
| *Acinetobacter baumannii*<br>17978VU *astNp*-30 bp-*lux* | This manuscript | LP948 |
| *Acinetobacter baumannii*<br>17978VU *astNp*-100 bp-*lux* | This manuscript | LP952 |

### Plasmid construction and mutant generation

All plasmids and oligonucleotides used in this manuscript are in Tables 2-3. DNA was amplified using Q5 High Fidelity 2X Master Mix (NEB) or Green GoTaq Master Mix (Promega). To generate a deletion mutant of *astR*, the pFLP2 vector with carbenicillin resistance and sucrose counterselection was used for allelic exchange^52^. Primers AW5 and AW6 were used to amplify 1000 bp downstream of the 3’ flanking region of *astR* and primers AW7 and AW8 were used to amplify 1000 bp upstream of the 5’ flanking region. The FRT flanked kanamycin resistance cassette was amplified using primers LP154 and LP155 from pKD4^53^. The pFLP2 backbone was digested using BamHI and KpnI restriction enzymes (NEB). Hifi assembly mix (NEB) was used to clone the kanamycin cassette, upstream, downstream regions into the digested pFLP2 backbone and amplified in *E. coli* DH5α λ*pir116*. Plasmid constructs were transformed into *A. baumannii* via electroporation, selected for Kn^R^ colonies, and colonies were further screened for Carb^R^ and sucrose^S^. PCR-confirmed merodiploids were plated to 10% sucrose to select for loss of pFLP2. Sucrose^R^ colonies were screened for Kn^R^ and Carb^S^. The knockout mutant was confirmed via PCR and whole genome sequencing (SeqCoast Genomics). To complement *astR* in the chromosome, *astRp* and *astR* were amplified using primers DB3 and DB4, and pKNOCK-mTn*7*^54^ was digested using BamHI and KpnI restriction enzymes (NEB), and the resulting DNA fragments were assembled with Hifi assembly mix (NEB). The pKNOCK-mTn*7* constructs were conjugated and transposed into the *A. baumannii* chromosome using the four-parent mating method described by Carruthers *et al.*^54^. To generate the *A. baylyi* Δ*astR*::Kn strain, pLDP311(pFLP2-*A. baylyi ΔastR*::Kn) was generated similarly to described above, digested with BamHI/KpnI (NEB), the resulting band with homology regions was gel extracted, and 1 μg linear DNA was added to 70 μL of *A. baylyi* culture and incubated for 3-6 h at 30°C with shaking at 250 rpm, then streaked to LB Kn to select for double crossovers, as described^55^, and confirmed by PCR. pWH1266 was digested with BamHI and KpnI to generate *A. baumannii* and *A. baylyi astR* complementation vectors with inserts from primers JG128-131 and JG149-152.

**Table 2:** Plasmids.

| Identifier | Description | Source |
| --- | --- | --- |
| pFLP2 | CarbR; allelic exchange vector with sucrose resistance | (Hoang *et al.*, 1998)<sup>52</sup> |
| pRK2013 | KnR; mobilization helper plasmid | (Figurski and Helinski, 1979)<sup>67</sup> |
| pKD4 | FRT-flanked kanamycin resistance cassette | (Datsenko and Wanner, 2000)<sup>53</sup> |
| pWH1266 | CarbR; shuttle plasmid for *E. coli* and *Acinetobacter* | (Hunger *et al.*, 1990)<sup>68</sup> |
| pKNOCK-mTn7-Amp | mini-Tn7-AmpR on a suicide vector containing the R6K γ-ori | (Carruthers *et al.*, 2013)<sup>54</sup> |
| pET15b | N-terminal his<sub>6</sub>-tag with T7 inducible promoter, three cloning sites, and AmpR | (Novagen, EMD Millipore, Darmstadt, Germany) |
| pLDP12 | pMU368(Tet)-no promoter-*lux* | (Juttukonda *et al.*, 2016)<sup>56</sup> |
| pLDP87 | pMU368(Tet)-no promoter-*lux-2* | This manuscript |
| pLDP90 | pFLP2-Δ*astR*::Kn | This manuscript |
| pLDP92 | pLDP87-*astNp-lux* | This manuscript |
| pLDP103 | pKNOCK-mTn7-*astRp-astR* | This manuscript |
| pLDP108 | pLDP87-*astRp-lux* | This manuscript |
| pLDP109 | pKNOCK-mTn7-*astRp*-His<sub>6</sub>-*astR* | This manuscript |
| pLDP163 | pLDP87-*astPp-lux* | This manuscript |
| pLDP208 | pET15b-His<sub>6</sub>-*astR* | This manuscript |
| pLDP265 | pLDP87-*astNp*-30 bp-*lux* | This manuscript |
| pLDP267 | pLDP87-*astNp*-75 bp-*lux* | This manuscript |
| pLDP268 | pLDP87-*astNp*-100 bp-*lux* | This manuscript |
| pLDP289 | pLDP87-*ACX60_10810p-lux* | This manuscript |
| pLDP311 | pFLP2-*A. baylyi* Δ*astR*::Kn | This manuscript |
| pLDP336 | pWH1266-*astRpA. baylyi-astRA. baylyi* | This manuscript |
| pLDP337 | pWH1266-*astRpA. baumannii-astRA. baumannii* | This manuscript |
| pLDP344 | pWH1266-*astRpA. baylyi-astRA. baumannii* | This manuscript |
| pLDP345 | pWH1266-*astRpA. baumannii-astRA. baylyi* | This manuscript |

**Table 3:**
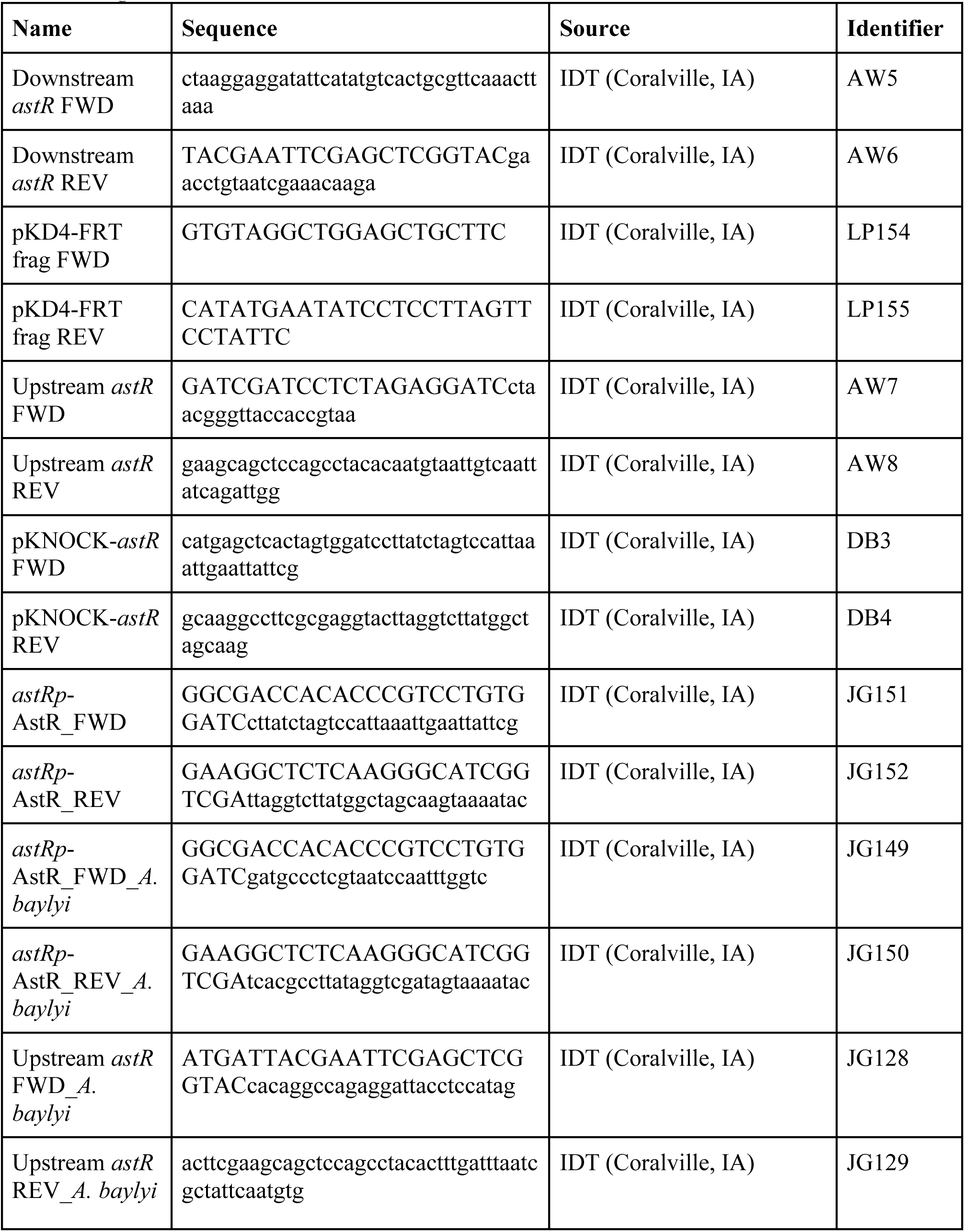
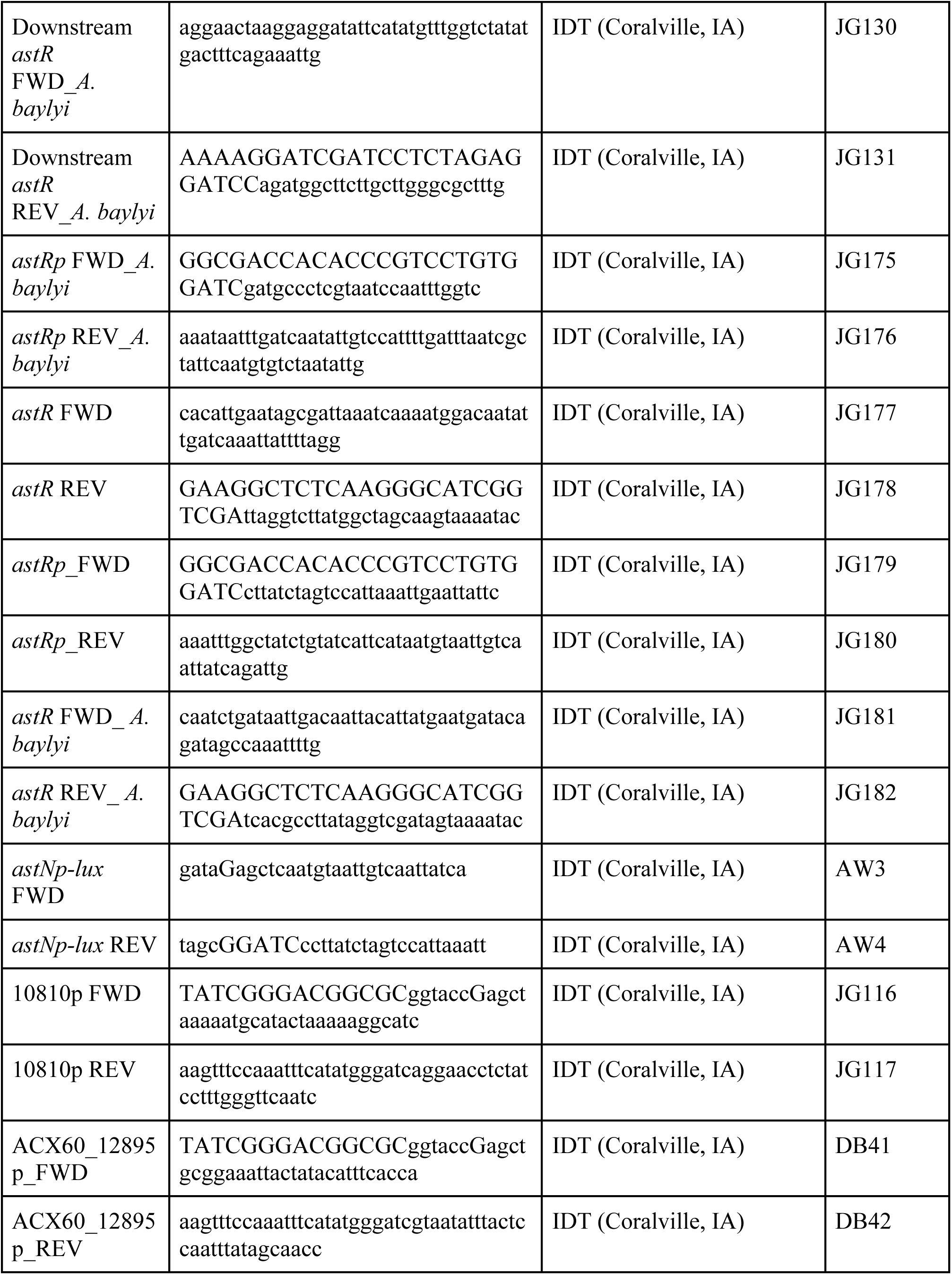
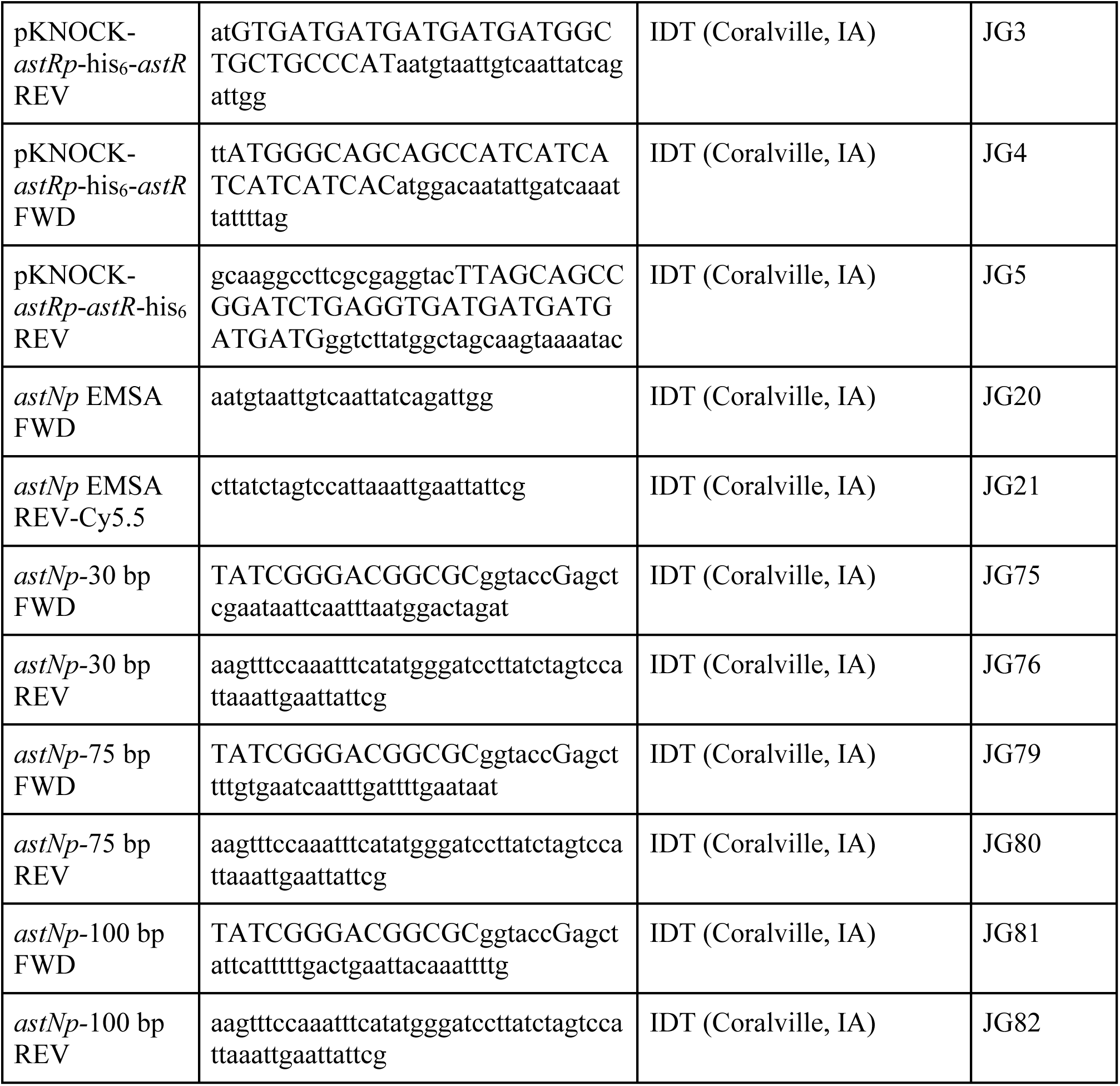
Oligonucleotides.

The pLDP87 *lux* reporter plasmid was derived from pMU368-*tet-lux*^56^. Sequencing of pMU368-*tet*-*lux* found 28 basepairs of DNA from *Streptococcus pneumoniae* strain Xen35 immediately upstream the RBS for *luxA*. This exogenous DNA was digested out of pMU368-tet-lux using BamHI and SpeI, and the lux operon was amplified using primers LP395 and LP396 and re-cloned to generate pLDP87 (pMU368-*lux-2*). Strain genotypes were analyzed via Sanger sequencing (UIC Genomic Research Core) or whole genome sequencing (SeqCoast Genomics).

### AstR phylogenetic tree and pairwise sequence alignments

Phylogenetic analyses of AstR were performed using RAxML v8.2.12 (Randomized Axelerated Maximum Likelihood)^55^. Homologs of AstR were identified **with** OrthoFinder v2.5.5^56,57^ from 126 proteomes spanning the genera *Acinetobacter*, *Pseudomonas*, *Enterobacter*, *Klebsiella*, and *Escherichia*, and the tree was rooted with *Nitrosomonas* species from the β-proteobacteria. The identified orthologs represent proteins descended from the last common ancestor of this group. Protein sequences were aligned with MUSCLE v5.3 (osx-arm64 build)^58^ under default settings, and the resulting multiple sequence alignments were analyzed with RAxML. To determine the most appropriate amino acid substitution model, RAxML compared likelihoods across models and selected the LG model^59^ as best-fitting. Statistical support was assessed using 100 rapid bootstrap replicates, seed=12345. The resulting phylogenetic tree was visualized, rooted with the *Yersinia enterolitica* AsnC protein (WP_005161192.1) and annotated with iTOL v6 (Interactive Tree Of Life)^57^. The species and AstR ortholog protein IDs used to construct the RAxML tree are listed in Table 4.

**Table 4:**
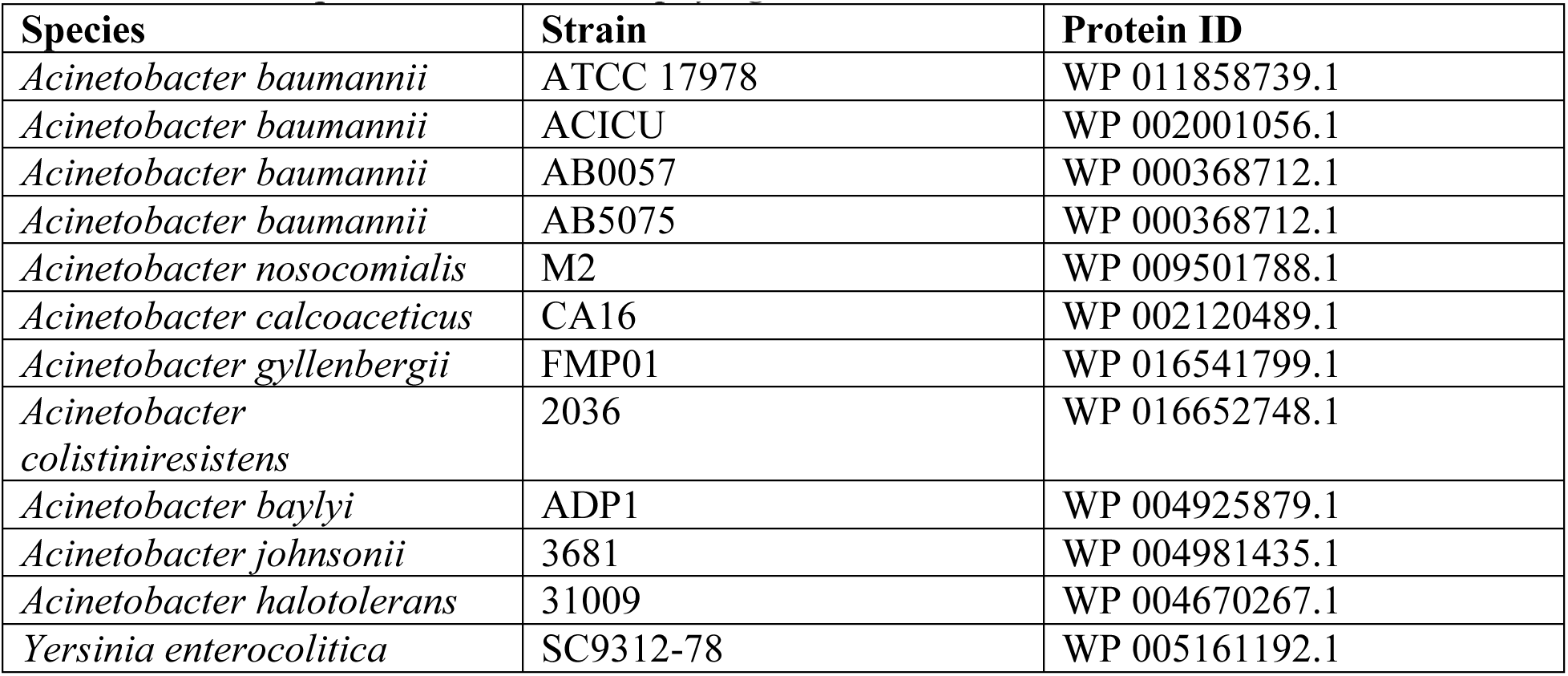
Protein sequences used in AstR phylogenetic tree.

RAxML performs a search of the space of possible phylogenetic trees using maximum likelihood. This search involves randomly perturbing the randomly-generated starting tree, evaluating the likelihood of each perturbed tree, and selecting the tree with the highest likelihood as the new starting point for the next round of perturbation. To assess the statistical support for the inferred tree, bootstrap analysis was performed, in which the original alignment is randomly resampled with replacement to generate 100 bootstrap replicates. Each bootstrap replicate was then analyzed using the same tree search procedure described above. The bootstrap support value for each branch of the tree is calculated based on the frequency with which that branch appears in the bootstrap trees.

Pairwise alignments of *A. baumannii* ATCC 17978VU and *A. baylyi* ATCC ADP1 AstR protein sequences were performed using EMBOSS Needle pairwise sequence alignment^58^.

### Bacterial growth and luminescence assays

Strains of *A. baumannii* and *A. baylyi* were grown at 37°C with 180 rpm shaking overnight in 3 mL LB with appropriate antibiotics for strains with plasmids. Overnight cultures were diluted 1:100 in PBS and then 1:100 into 99 μL of minimal media (10,000-fold total dilution) in 96-well flat bottom plates. Bacterial density by optical density at 600 nm (OD_600_) and luminescence were quantified in a Biotek Synergy plate reader at 37°C with shaking. All growth in minimal media for growth experiments contained 16.5 mM concentrations of carbon and 18.6 mM of nitrogen sources, added to carbon and nitrogen-free M9 medium^59^ with 0.1X Vishniac trace minerals^60^. For luminescence reporter assays, *A. baumannii* cultures bearing a pMU368-*astNp-lux,* pMU368-truncated-*astNp-lux*, pMU368-synthetic *astNp-lux* or pMU368-*astRp-lux* were grown. The luminescence readings were normalized to OD_600_ at ∼0.4 or as indicated.

### RNA Sequencing

The *A. baumannii* ATCC 17978VU WT and Δ*astR* mutant were grown overnight at 37°C in LB broth and then subcultured 1:1000 into fresh LB. After 3.5 h of growth at 37°C with shaking, cells were centrifuged at 4,137 x *g* for 5 minutes and washed twice with no-nitrogen 1X M9 salts^59^. Samples were resuspended in 1 mL of the 1X M9 salts with No Nitrogen and added to 9 mL of M9 media^59^ containing 16.5 mM concentrations of carbon sources (ornithine or succinate) and 18.6 mM of nitrogen sources (ornithine or NH_4_), as well as 0.1X Vishniac trace minerals^60^. The resuspended cells were incubated in the media for 1 h at 37°C. Cells were harvested by centrifugation at 4,137 x *g* for 10 min. Supernatant was discarded and residual liquid was pipetted out from the pellets. Pellets were stored at -80°C until RNA extraction. Pellets were thawed on ice and resuspended in 100 µL of RNAse free TE Buffer pH 8.0. The samples were lysed and homogenized in a Yellow Bullet Blender tube (MidSci) at Bullet Blender (NextAdvance) speed 8 for 3 minutes. The Qiagen RNeasy was used to isolate and purify the RNA from the samples, following the manufacturer’s protocol. Additional sample preparation and sequencing were performed by SeqCoast Genomics. Samples were prepared for sequencing using an Illumina Stranded Total RNA Prep Ligation with Ribo-Zero Plus Microbiome and unique dual indexes. Sequencing was performed on the Illumina NextSeq2000 platform using a 300 cycle flow cell kit to produce 2×150 bp paired reads. Read demultiplexing, read trimming, and run analytics were performed using DRAGEN v3.10.12, an on-board analysis software on the NextSeq2000. Reads were mapped to the *A. baumannii* ATCC 17978VU reference genome (CP012004.1, CP049364.1, CP049365.1, CP012005.1)^61,62^ and analyzed using DESeq2 1.44.0^63^. Because DESeq2 calculates the log_2_fold change using the baseMean across all samples analyzed, there was a -log_2_fold change reported for the *astR* transcript above the cutoff threshold in Δ*astR* samples. The sequencing data were manually inspected and confirmed that the *astR* transcript was not present Δ*astR* samples. Thus, *astR* log_2_fold change data were removed from Δ*astR* comparisons. For genes with expression fold changes with absolute value >4 and adjusted *p* value < 0.05, pathway analysis was performed using the Kyoto Encyclopedia of Genes and Genomes (KEGG)^64,65^.

### Cloning, expression, and purification of recombinant AstR

The ORF of *astR* was cloned into pET15b to acquire an N-terminal His_6_ tagged construct. The *astR* ORF was isolated with the primer pair JG41 and JG42, and ligated into digested pET15b (NdeI, XhoI) with HiFi DNA Assembly kit (NEB) to generate pLDP208(pET15b-*astR*) which was amplified in *E. coli* DH5α *pir116*. *E. coli* BL21 (DE3) RIL was transformed with pLDP208 and grown in terrific broth (tryptone 12 g/L, yeast extract 24 g/L, K_2_HPO_4_ 9.4 g/L, KH_2_PO_4_ 2.2 g/L, 0.8% glycerol, MilliporeSigma) with carbenicillin 75 mg/L at 37°C to an OD_600_ of 0.6 before induction with 0.1 mM IPTG and incubated at 15°C at 250 rpm for 16 h. Cells were harvested by centrifugation at 4,137 x *g* for 10 minutes and resuspended in B-PER™ Bacterial Protein Extraction Reagent (Thermo Fisher). The lysing cells were incubated at room temperature for 1 h with 2D rotor shaking. Lysate was centrifuged at 16,000 x *g* for 5 minutes to pellet insoluble proteins and intact cells, and the supernatant was resuspended 1:1 in lysis buffer (50 mM NaH_2_PO_4_, 300 mM NaCl, 10 mM imidazole, pH 8.0) to generate cleared lysate. Ni-NTA agarose resin (Qiagen) was centrifuged at 4,137 x *g* for 5 minutes and resuspended four volumes of lysis buffer. The resin was centrifuged again and supernatant was discarded. The pelleted resin was resuspended in two volumes of cleared lysate, which was divided into four equal portions for rocking on a 2D rotor at 4°C for 1 hour. The resuspension was centrifuged to pellet resin bound protein, all but 5 mL of the supernatant was decanted, the pellet resuspended in the remaining 5 mL supernatant, and loaded into four Poly-Prep Chromatography Columns (Bio-Rad). Flowthrough was collected and the column was washed with five bed volumes of wash buffer (50 mM NaH_2_PO_4_, 300 mM NaCl, 50 mM imidazole, pH 8.0), two times. His_6_-AstR was eluted in one bed volume elution buffer (50 mM NaH_2_PO_4_, 300 mM NaCl, 500 mM imidazole), four times. Elution fractions with pure His_6_-AstR were subjected to buffer exchange in PD-10 desalting columns (Cytiva) to the EMSA buffer (20 mM Tris HCl, 2.5 mM MgCl_2_, 0.45 mM EDTA, 50 mM NaCl, 10% glycerol, pH 7.5), Size Exclusion Buffer (50 mM Tris, 300 mM NaCl, pH 8), or Mass Spec Buffer (50 mM ammonium acetate, pH 7). His_6_-AstR was concentrated using an Amicon Ultra 3K MWCO Centrifugal filter device (MilliporeSigma) at 4,000 x g for 10 minutes. His_6_-AstR was stored at -80°C. Depiction of promoter features was based on literature for the +1 transcription start site and PhiSite Promoter Hunter with default *E. coli* matrix for -10 and -35 RNA polymerase binding sites^32,66^.

### Electrophoretic mobility shift assays

Promoter DNA was amplified by PCR using primer pairs in which the reverse primer had a 5’-Cy5.5 fluorophore tag (JG20 and JG21 for *astNp*) (Integrated DNA Technologies,) and purified using a Qiagen PCR purification kit (Qiagen). 5 nM of purified fluorescent promoter was incubated at room temperature in the dark for 30 minutes with His_6_-AstR at concentrations of 0.5-10 µM in 1X EMSA buffer (final concentrations: 20 mM Tris HCl, 2.5 mM MgCl_2_, 0.45 mM EDTA, 50 mM NaCl, 10% glycerol, and 5 mM dithiothreitol added fresh). EMSA buffer was prepared as a 2X stock and diluted appropriately to ensure appropriate final concentrations with the addition of protein, DNA, and 5 mM additional molecules as indicated. The 7.5% Mini-PROTEAN® TGX™ Gel (Bio-Rad) was prerun in Tris-buffered EDTA buffer (89 mM Tris, 89 mM boric acid, 2 mM EDTA, pH 8.3) (Bio-Rad) on ice at 100 V for 30 minutes. The incubated samples were then separated by gel electrophoresis at room temperature. Gels were stained with SYBR green (Invitrogen) diluted 1:10,000 in TBE for 10 min in the dark, washed twice with H_2_O, and visualized with a Chemidocgel imager (Bio-Rad, Hercules, CA) for Cy5.5 fluorescence.

### Mouse experiments

*Conventional mice.* Six-week old Swiss Webster mice were purchased from Charles River Lab. These mice were fed a diet of autoclaved 5L79 (PMI Nutrition International 5L79) ensured to maintain appropriate vitamin levels. Mice were housed in the University of Illinois Chicago Biological Resources Laboratory (BRL). Once delivered, mice were adapted to the BRL facility for one week. Then, 1 g/L gentamicin water was administered to the mice for a five-day (day -7 through -2) pretreatment, with replacing the antibiotic water every two to three days. Normal drinking water was administered to mice two days (-2) before gut colonization. *A. baumannii* 17978VU strains WT *att*::mTn*7*(carb^R^) and the Δ*astR*::Kn mutant were grown at 37 °C for 16 hours in 10 mL of LB Broth. Next, cells were centrifuged at 4,000 x *g*, 4 °C for 7 minutes, washed two times with PBS, resuspended and normalized in PBS to 1 × 10^10^ colony forming units (CFU)/mL. On day 0, mice were orogastrically gavaged with 100 μL (1 ×10^9^ CFU) of a 1:1 mixture of *A. baumannii* 17978VU WT:Δ*astR* mutant. Mouse fecal pellets were collected on days 0, 1, 4, 7 and 9 days post inoculation. Mice were euthanized humanely 10 days post inoculation by CO_2_ asphyxiation and gastrointestinal organs were sterilely removed. To quantify *A. baumannii* CFU, mouse fecal pellets were weighed, resuspended in 1 mL of PBS, serial diluted to 10^−7^ and plated to the proper selective LB plates with antibiotics. CFU were enumerated from 50 mg/L carbenicillin and 5 mg/L chloramphenicol LB agar for *A. baumannii* 17978VU WT *att*::mTn*7* or 40 mg/L kanamycin sulfate and 5 mg/L chloramphenicol LB agar for *A. baumannii* Δ*astR*::Kn. Organs were homogenized in PBS with metal beads in a NextAdvance Bullet Blender tissue homogenizer speed 8 for 10 minutes. CFU from feces were reported per g of fecal sample, whereas CFU from an organ were reported per whole organ.

*Germ-free mice*. Germ-free C57BL/6J mice were bred in the BRL facility at the University of Illinois Chicago in a room with 14 h:10 h light/dark cycles and 70–76°F and 30%– 70% humidity. Mice were kept in isolators purchased from Park Bioservices LLC. Mice were fed autoclaved mouse diet 5L79 (PMI Nutrition International 5L79) and autoclaved super Q water *ad libitum* and were 8–12 weeks of age. Germ-free condition was tested at least once a month with aerobic liquid cultures (brain heart infusion, BHI), solid cultures (blood agar plates, Thermo Sci Remel), fungal cultures (Sabouraud slants), anaerobic liquid (BHI) and solid cultures (*Brucella* agar, Thermo Sci Remel) from isolators’ swabs, fecal samples, and fungal traps placed inside the isolators. Fecal samples were also tested with Gram staining and qPCR to detect bacterial DNA.

*Ethics and approvals*. Animal care protocols were approved by the UIC Institutional Animal Care and Use Committee (protocol numbers 23-119 and 22–192) according to the Animal Care Policies at UIC, the Animal Welfare Act, the National Institute of Health and the American Veterinary Medical Association. For experimental endpoints, animals were humanely euthanized in accordance with American Veterinary Medical Association (AVMA) guidelines. No statistical methods were used to pre-determine sample sizes.

### Fecal slurry experiments

To generate an *ex vivo* model of the gut environment, a fecal slurry was created using 25-30 mg of a single mouse fecal pellet resuspended in 1 mL PBS. In 96-well flat bottom plates, 0.1 mL fecal slurry were inoculated with 1 uL of the indicated strain. The plates were incubated at 37°C with shaking and luminescence was monitored in a Biotek Synergy plate reader. Growth in fecal slurry was confirmed via CFU enumeration on 10 mg/L chloramphenicol plates to select for *A. baumannii*. For fecal slurry assays, feces were collected from conventional Swiss Webster mice or germ-free C57BL/6 mice fed autoclaved 5L79.

### Statistical analysis

Each measurement was taken from a distinct biological sample (*e.g.* an individual mouse or bacterial culture from a single colony). Statistical analyses were performed with Microsoft Excel 16.77.1, GraphPad Prism 10.5.0, and DESeq2 1.44.0 for RNA-seq analysis. Values below the limit of detection are graphed at the limit of detection for statistical purposes. The respective statistical tests are described in the figure legends. All *p* values for Student’s *t* tests, Mann-Whitney tests, and Mantel-Cox tests are two-sided. Figures were prepared in Adobe Illustrator 28.7.9.

## DATA AVAILABILITY STATEMENT

RNA-seq sequencing files will be deposited to the National Center for Biotechnology Information (NCBI) Gene Expression Omnibus (GEO) database prior to publication. All data used to generate graphs will be uploaded to Zenodo prior to publication.

## CODE AVAILABILITY STATEMENT

Code for obtaining data, running OrthoFinder and generating phylogenetic trees can be found at https://github.com/JonWinkelman/astR_trees_geary2025.

## ACKNOWLEDGMENTS

We thank the Palmer laboratory for helpful comments. We thank Anita Waye for generating pLDP90, Jared Winkelman at Trestle, LLC, for assistance with RNA-seq bioinformatics analysis, Jordan J. Jesse for assistance with *astPp* reporter strains, William Rowley for assistance with AstR mass spectrometry, and Ciera Duffy, Edward Eshoo, and Judith Behnsen for providing germ-free mice fecal samples. This work was supported by startup funds from the University of Illinois Chicago to M.H. and L.D.P., Burroughs Wellcome Fund PATH Award 1019120.01 to F.A., Pioneer in Research Award from the Michael Reese Research & Education Foundation to L.D.P., and the National Institutes of Health awards R01AI175223 to Judith Behnsen, R01AI120994 to F.A., and R01AI189516 and R01AI192538 to L.D.P. The content is solely the responsibility of the authors and does not necessarily represent the official views of the National Institutes of Health or other funders.

## AUTHOR CONTRIBUTIONS

Conceptualization, J.G.H. and L.D.P. Methodology, J.G.H., X.R., N.P., I.C.A., F.A.3^rd^, M.T.H., and L.D.P.; Investigation, J.G.H., X.R., D.A.B., N.T.T.P., I.C.A., B.K., and J.D.W; Writing—Original Draft, J.G.H., X.R., and L.D.P.; Writing—Review & Editing, J.G.H., X.R., D.A.B., N.T.T.P., I.C.A., B.K, J.D.W., F.A.3^rd^, M.T.H., Visualization, J.G.H., X.R., N.T.T.P., M.T.H., L.D.P.; Supervision, F.A.3^rd^, M.T.H., L.D.P. Funding Acquisition, F.A.3^rd^, M.T.H., L.D.P.

