## Supplementary information for "A regulator of amino acid catabolism controls *Acinetobacter baumannii* gut colonization"

**A**

```
A. baumannii AstR 1   MDNIDQIILGLLKDNARISVTELAEKVHVSRATIKKRMEYLESSGIITGY  50
                      |::.|..|||||:||||:|||:||.||.||||||:||:||:|:.||||||
A. baylyi AstR    1   MNDTDSQILGLLRDNARMSVTDLAAKVKVSRATIQKRIEYMENQGIITGY  50
                       +                    +     ++  + + +            +

A. baumannii AstR 51  TVRFKPNAERNVIRAWMSIMVEGTKAQSVIKELRLESAVECLHKTNGKWD  100
                      |||.|||:|:|:|||||:|||||.||.:||:|||::.||..||.||||||
A. baylyi AstR    51  TVRIKPNSEQNMIRAWMNIMVEGNKASAVIRELRIDPAVHTLHTTNGKWD  100
                         +    +     +           +     +          ## # ##
                                                                +

A. baumannii AstR 101 ILVELRSDTLENFDKVLERIRNISGIYNSETSILLASHKT  140
                      :||||:|:.|.:|||:|||||.|:||||:||||||:::|.
A. baylyi AstR    101 LLVELKSENLASFDKILERIRTINGIYNTETSILLSTYKA  140
                          +     +     #   #       ###
```

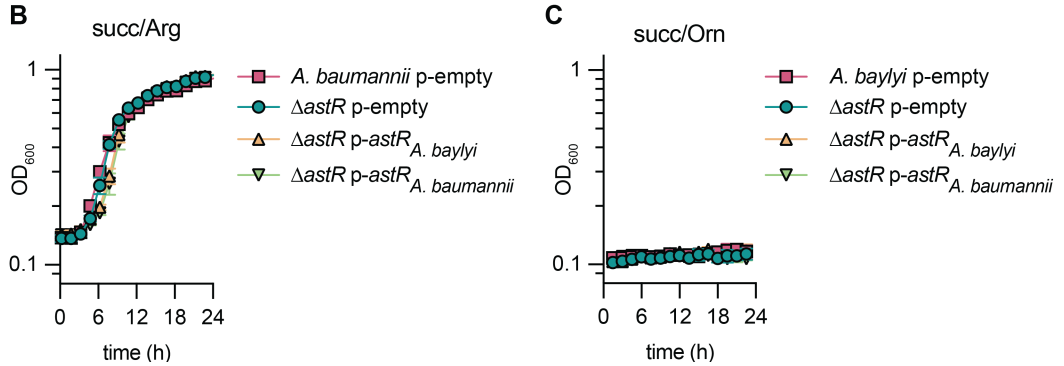


**Figure S1: Additional data of *A. baumannii* and *A. baylyi* AstR.**
(**A**) Protein sequence alignment of AstR encoded by *A. baumannii* ATCC 17978 and *A. baylyi* ATCC ADP1. The amino acid sequence identity is 96/140 (68.6%) and similarity is 123/140 (87.9%) by EMBOSS Needle pairwise sequence alignment[1]. Below the sequence, important residues from *E. coli* AsnC are marked with # for ligand binding and + for DNA binding[2].
**(B-C)** *A. baumannii* ATCC 17978VU or *A. baylyi* ATCC ADP1 wildtype and Δ*astR* strains with empty or *astR* complementation introduced in the pWH1266 plasmid. The pWH1266 plasmids used endogenous the *astR* promoter from the host strain. Strains were grown in nitrogen-free M9 minimal media with succinate at 16.5 mM and arginine or ornithine at 18.6 mM, as indicated (n = 3; mean ± SD, experiments were performed at least twice with similar results).

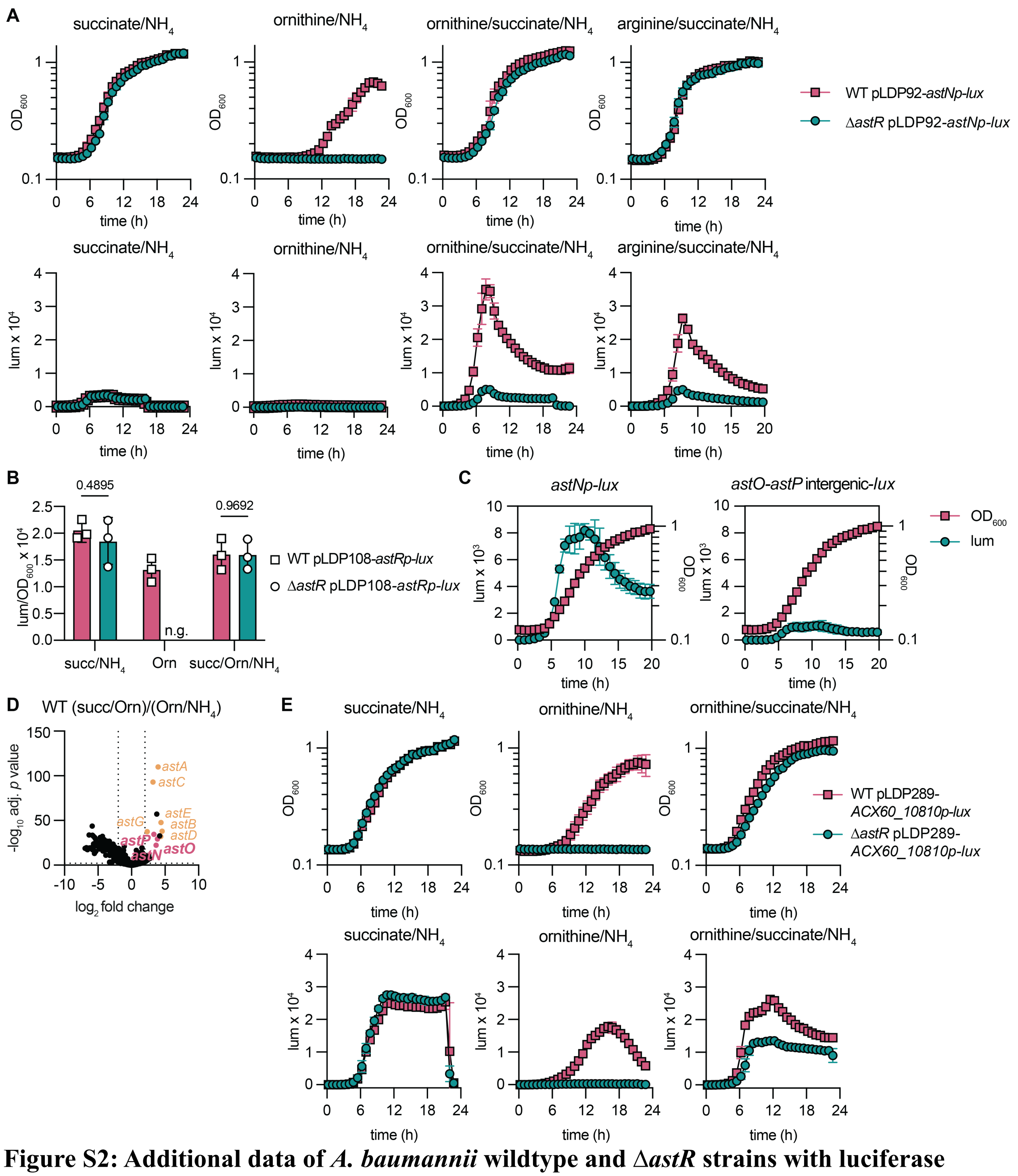


**Figure S2: Additional data of *A. baumannii* wildtype and Δ*astR* strains with luciferase reporters of *astNOP*, *astR*, and *ACX60_10810* promoter activity.**

**(A)** *A. baumannii* ATCC 17978 wildtype (WT) and Δ*astR* strains containing an *astN* promoter (*astNp*) driving luciferase expression from the *lux* operon were grown in nitrogen-free M9 minimal media minimal media with succinate (succ), ornithine (Orn), and arginine at 16.5 mM and $NH_4Cl$ at 18.5 mM, as indicated (n=3; mean ± SD, experiments were performed at least twice with similar results).

**(B)** *A. baumannii* ATCC 17978 wildtype (WT) and Δ*astR* strains containing an *astN* promoter (*astNp*) driving luciferase expression from the *lux* operon were grown in nitrogen-free M9 minimal media minimal media with succinate (succ), ornithine (Orn), and arginine at 16.5 mM and $NH_4Cl$ at 18.5 mM, as indicated. Luminescence was normalized to optical density at 600 nm ($OD_{600}$) at 0.25. N.g. indicates no growth in that condition (n = 3; mean ± SD; *p* by multiple unpaired *t*-tests; experiments were performed at least twice with similar results).
**(C)** *A. baumannii* ATCC 17978 wildtype containing an *astP* promoter (*astPp*) driving luciferase expression from the *lux* operon were grown in nitrogen-free M9 minimal media minimal media with succinate (16.5 mM) and ornithine (18.6 mM) (n = 3; mean ± SD, experiments were performed at least twice with similar results).
**(D)** RNA-seq of *A. baumannii* ATCC 17978 wildtype (WT) incubated for 1 h in M9 minimal medium with succinate (succ; 16.5 mM) and ornithine (18.6 mM) or ornithine (16.5 mM) and NH4Cl (18.6 mM) as the sole carbon and nitrogen sources, respectively. Significantly changed *ast* genes are colored and labeled. Each dot is one gene (n = 3; adj. *p* value and fold-change were determined by DESeq2; dotted lines shown at significance cut-offs |log2 fold change|=2 and adj. *p* value=0.05).
**(E)** *A. baumannii* ATCC 17978 wildtype (WT) and Δ*astR* strains containing an *ACX60_10810* promoter (*ACX60_10810p*) driving luciferase expression from the *lux* operon were grown in nitrogen-free M9 minimal media minimal media with succinate (succ) and ornithine (Orn) at 16.5 mM and $NH_4Cl$ at 18.6 mM, as indicated (n = 3; mean ± SD, experiments were performed at least twice with similar results).
$OD_{600}$, optical density at 600 nm; WT, wildtype; *astNp, astNOP* operon promoter; lum, luminescence; *astRp*, *astR* promoter; succ, succinate; Orn, ornithine; *10810p*, *ACX60_10810* promoter.

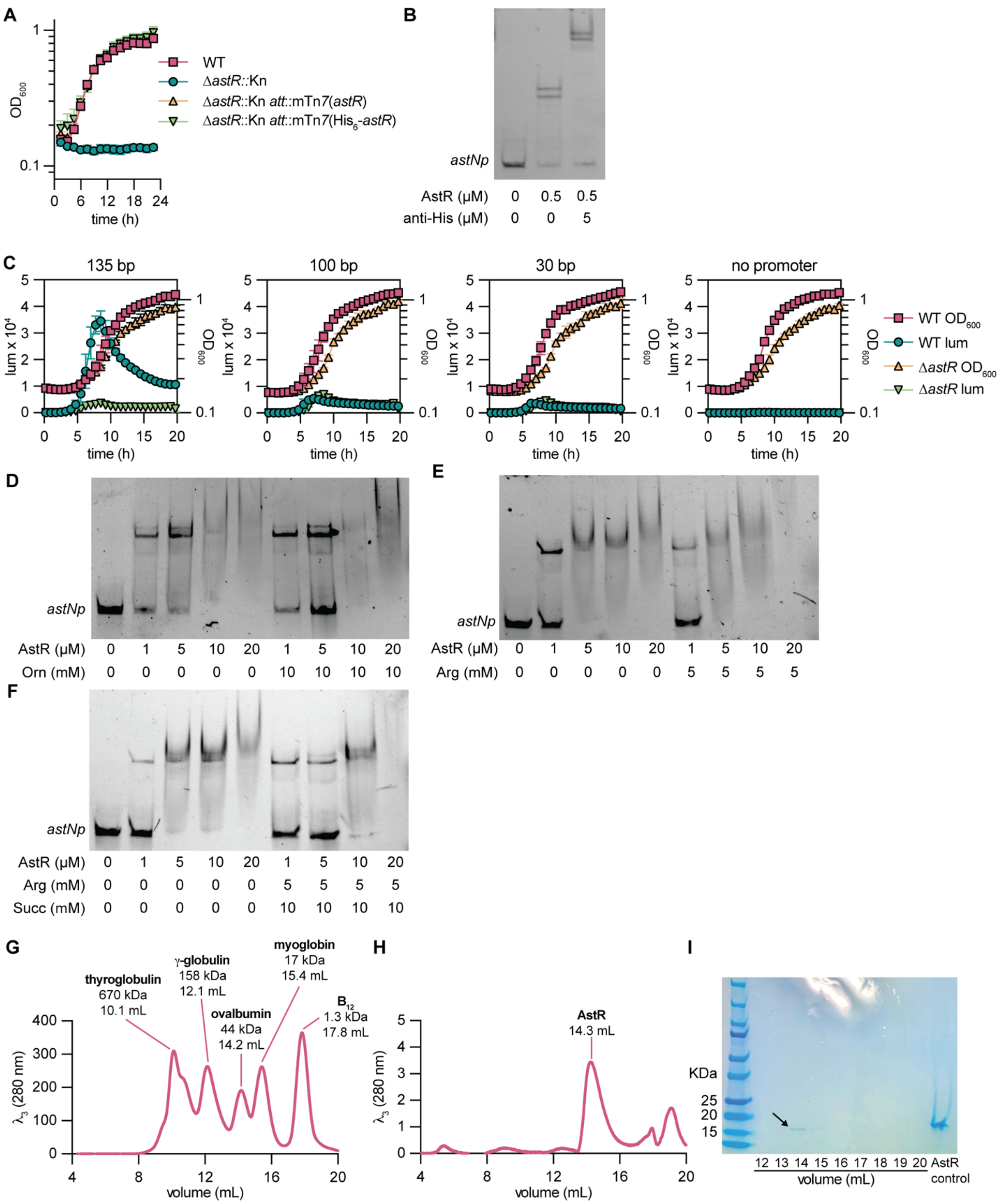


**Figure S3: Additional data for His6-AstR function and oligomerization state.**

**(A)** *A. baumannii* ATCC 17978 wildtype (WT) and Δ*astR* strains containing mTn*7* with wildtype or His$_6$-AstR were grown in nitrogen-free M9 minimal medium with ornithine (16.5 mM) as the sole carbon and nitrogen source (n = 3, mean ± SD, experiments were performed at least twice with similar results).

**(B)** His$_6$-AstR, Cy5.5-labeled *astNp*, and anti-His antibody were incubated together at the indicated concentrations for electrophoretic mobility shift assays (EMSA).

**(C)** The 135 bp, 100 bp, and 30 bp intergenic regions upstream of *astN* were cloned upstream of a *lux* luciferase reporter or with a no promoter control. WT and Δ*astR* strains with the *lux* reporters were incubated in nitrogen-free M9 minimal medium with ornithine at 16.5 mM, succinate at 16.5 mM, and $NH_4Cl$ at 18.6 mM. ($n = 3$; mean ± SD, experiments were performed at least twice with similar results).

**(D-F)** His$_6$-AstR, Cy5.5-labeled *astNp*, and indicated metabolites were incubated together at the indicated concentrations for EMSA.

**(G-H)** Size exclusion chromatography of standards and His$_6$-AstR.

**(I)** Polyacrylamide gel of size exclusion fractions stained with SimplyBlue confirm the peak at 14 mL is likely AstR.

$OD_{600}$, optical density at 600 nm; mTn*7*, mini Tn*7*; *astNp, astNOP* operon promoter; lum, luminescence; Orn, ornithine.

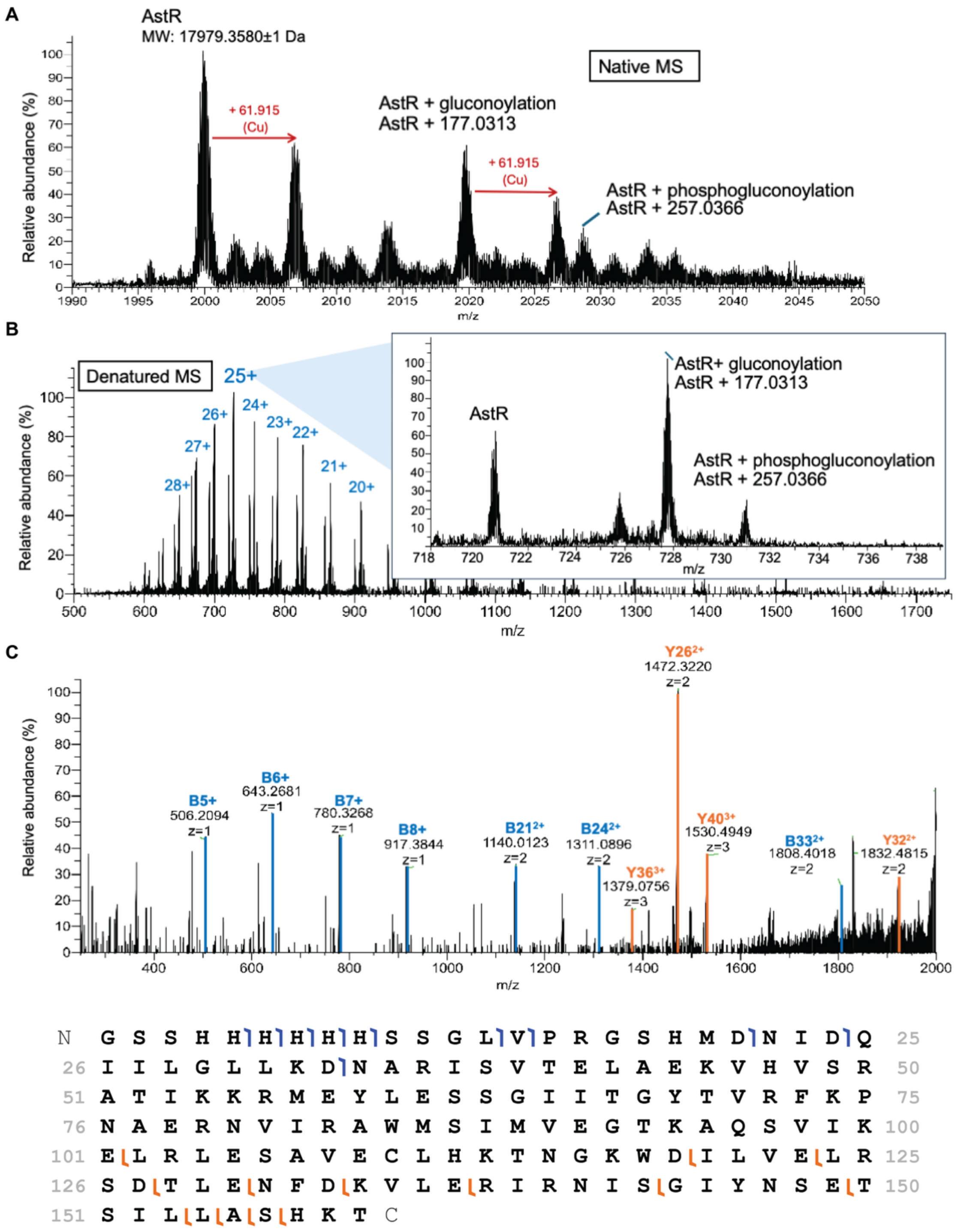


**Figure S4: AstR does not appear to bind any ligand as determined by native and denatured mass spectrometry.**

**(A)** Native mass spectrometry (MS) showed a complex spectrum indicating many distinct states of AstR. AstR appeared with the removal of N-formylmethionine. It was futher modified with α-

N-gluconoylation (+177.0313±1. Da) and α-N-6-phosphogluconoylation (+257.0366±1Da), supported by Geoghegan *et al.*, 1998[3]. Both AstR and modified AstR potentially bound to copper.

**(B)** Denatured LC-MS showed similar features of modified AstR.

**(C)** Tandem mass spectrum (MS2) of AstR with annotations of a few highly abundant b and y ions, which were detected and mapped by ProSight Lite[4]. Data representative of analysis from two independently purified protein preparations.

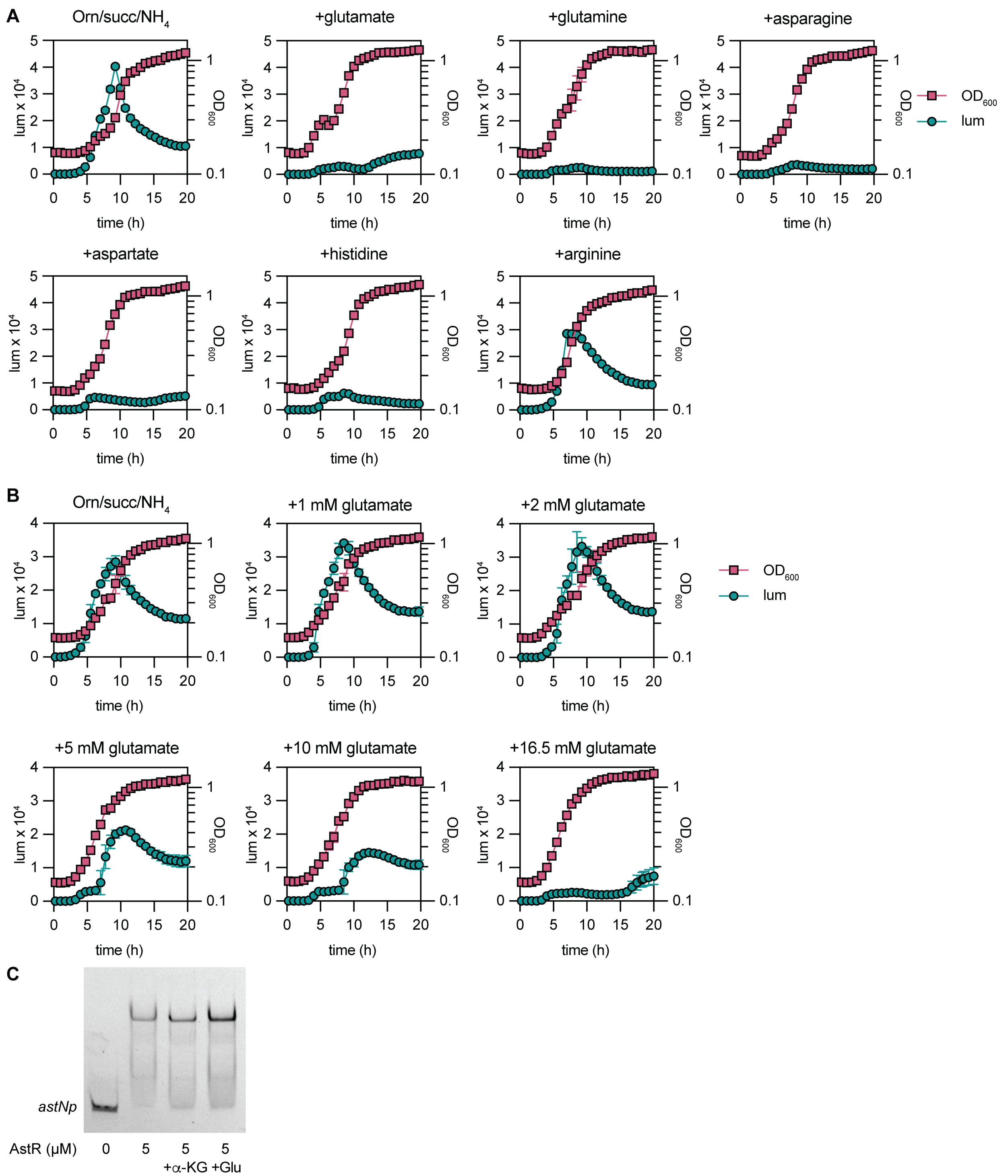


**Figure S5: Growth, luminescence, and EMSA in the presence of preferred amino acid carbon sources such as glutamate.**
**(A-B)** *A. baumannii* ATCC 17978 wildtype containing an *astN* promoter (*astNp*) driving luciferase expression from the *lux* operon was grown in nitrogen-free M9 minimal media minimal media with succinate (succ) and ornithine (Orn) at 16.5 mM and $NH_4Cl$ at 18.5 mM. In (A), 16.5 mM additional amino acids were added as indicated. In (B), glutamate was added at the indicated concentrations.

Luminescence and optical density at 600 nm ($OD_{600}$) are shown (n = 3, mean ± SD, *p* by one-way ANOVA with Dunnett's multiple comparisons to no additional amino acids; experiments were performed at least twice with similar results)

**(C)** Cy5.5-labeled *astNp* DNA was incubated with $His_6$-AstR at the indicated concentrations and alpha-ketoglutarate or glutamate at 5 mM for EMSA.

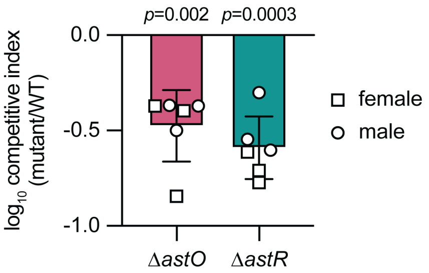


**Figure 6: Competitive index of *A. baumannii* ATCC 17978 wildtype (WT) compared to Δ*astO* and Δ*astR* for growth in fecal slurries from conventional female and male mice.** n = 6 (fecal samples from n = 3 female mice and n = 3 male mice); mean ± SD; *p* by one-sample *t* test).

### SUPPLEMENTAL METHODS

#### Size exclusion chromatography

Purified $His_6$-AstR was applied to a BioRad ENrich size exclusion chromatography (SEC) 650 10 × 300 mm column connected to a BioRad NGC medium pressure chromatography system and the BioRad ChromLab software was used for analysis. The column is stored in 20% ethanol while not being used, and then washed with water before every use. The column was equilibrated with 2.5 column volumes of the SEC buffer (50 mM Tris-HCl, 300 mM NaCl pH=8.0). Following equilibration, 200 µL of the sample was injected into the valve. Elution fractions were collected in 1-mL volumes from 4 mL to 26 mL. Fractions 9-17 (the fractions that exhibited peaks and those in between in ChromLab) were separated by on sodium dodecasulfate polyacrylamide gel electrophoresis (SDS-PAGE) and stained using SimplyBlue SafeStain (ThermoFisher). To estimate the size of the AstR oligomer, gel filtration standards containing a lyophilized mix of thyroglobulin, bovine γ-globulin, chicken ovalbumin, equine myoglobin, and vit $B_{12}$ (Bio-Rad, Hercules, CA) was passed through the column and their peaks were analyzed in ChromLab.

#### LC-MS, Native mass spectrometry acquisition and analysis

For native mass spectrometry (MS), purified $His_6$-AstR was exchanged into 50 mM ammonium acetate buffer, pH 7, using a PD-10 column (Cytiva). The sample was sterile filtered using 0.2 µm hydrophilic polyvinylidene fluoride (PVDF) syringe filters to remove large contamination. The system was first optimized with 0.1 mg/mL lysozyme solution. Ion spray parameters were set at 3000 V for spray voltage, 20 for sheath gas, 5 for aux gas, and 0 for sweep gas. About 2 µg was injected to the Orbitrap Exploris 120 (Thermo Fisher Scientific) with the flowrate of 10-20 µL/min through a low-flow metal needle. To acquire MS2 data for a specific *m/z* value, injection time was manually increased from 100 to 1000 ms to accumulate more ions. At the same time, the mass range was adjusted to collect ions of interest, decreased high-energy collisional dissociation (hcd) to 1 while collecting ions, and increased the number of microscans to 10 to verify the purity of ions in the HCD cell. Then, the HCD energy was gradually increased up to 35 for fragmentation.

For denatured analysis, the sample of $His_6$-AstR in 50 mM ammonium acetate buffer was injected into Vanquish HPLC connected to Obitrap Exploris 120 MS (Thermo Fisher Scientific). The protein was loaded on 250 mm monolith ProSwift™ RP-4H LC column with a flow rate of 0.150 µL/min. The mobile phase started with 95% water (with 0.1% formic acid) and gradually increased to 100% acetonitrile (with 0.1% formic acid). Data acquisition in LC-MS system was done by Xcalibur (Thermo Fisher Scientific). The spectra for both Native MS and LC-MS were averaged and analyzed by FreeStyle (Thermo Fisher Scientific).
